# Two high-quality rose genomes underpin a novel *Rosa* pangenome to advance rose genomics, phylogenetics, and breeding

**DOI:** 10.1101/2025.05.07.652600

**Authors:** Zijiang Yang, Elio G. W. M. Schijlen, M. Eric Schranz, Bas J. Zwaan, Dick de Ridder, Richard G.F. Visser, Paul F. P. Arens, Marinus J. M. Smulders, Robin van Velzen, Joost J. B. Keurentjes, Peter M. Bourke, Sandra Smit

## Abstract

*Rosa*, belonging to the family Rosaceae, encompasses more than 150 species which are widely distributed in the northern hemisphere. Renowned for their beauty, roses are cultivated throughout the world for ornamental purposes and the production of essential oils and perfumes. Despite their cultural and commercial significance, the genomic resources of wild *Rosa* species have not been studied comprehensively, hampering the understanding of their genetic diversity, evolutionary history, and breeding potential.

Here we report on high-quality *de novo* genomes for *Rosa sericea* and *Rosa rugosa*. By integrating these two *de novo* genomes with existing public genomic resources, we have built a Rosaceae panproteome and a *Rosa* pangenome (spanning wild, traditional garden, and modern rose lineages) using a De Bruijn graph (DBG)-based approach. A maximum likelihood (ML) phylogeny of 18 *Rosa* haplotypes based on 4,367 single-copy core homology groups (genes) provided robust evolutionary inference, confirming the basal position of *R. sericea*, and enabled a gene-based macrosynteny analysis across the pangenome. Our analyses revealed significant genomic diversity among species, extensive variation in core gene content, and lineage-specific transposable element (TE) expansion patterns that contribute to the variation in *Rosa* genome size and to species-specific adaptations. The pangenome also revealed biased diversification of homology groups potentially linked to phenotypic plasticity in *Rosa*. Specifically, our analysis of the rose scent-related gene family, *NUDX1*, uncovered its evolutionary trajectory in *Rosa*, in which TEs insertions provided putative novel regulatory elements that facilitated adaptive evolution in metabolic pathways.

This pangenomic study deepens our understanding of the genetic diversity and evolution of traits within the *Rosa* genus. In addition, the findings lay the foundation for future efforts to understand the genetic mechanisms driving trait evolution, which can support rose breeding.

## Background

Species of the genus *Rosa*, renowned for their ornamental value, are highly appreciated in modern horticulture. Driven by both aesthetic preferences and economic incentives, roses have been cultivated for thousands of years (Bendahmane et al., 2013). Based on its domestication history, the genus *Rosa* can be divided into two major groups: wild rose species, and cultivated roses, the latter including both traditional “Old garden roses” (e.g. *R. damascena* and *R. gallica*) developed during early cultivation, as well as modern roses (*R. hybrida*) that emerged from 18^th^ century hybridization between Chinese recurrent-blooming roses (e.g., *R. chinensis*) and European species (Wylie, 1954; Dubois et al., 2010; Bendahmane et al., 2013).

Natural hybridization is widespread among *Rosa* populations. The Caninae section exemplifies this phenomenon, where most species exhibit pentaploid somatic status (2n=5x=35) believed to derive from multiple ancient hybridization events (Ritz et al., 2005; Lim et al., 2005; Reichel et al., 2023). A recent genomic study also uncovered hybrids in diploid lineages, where natural crosses between *R. roxburghii* and *R. longicuspis* likely gave rise to *R. sterilis* (2n=2x=14) (Zong et al., 2024). The extensive reticulation driven by both natural and human-driven selection pressures has contributed to a remarkable phenotypic diversity in floral traits (e.g., petal color and number), growth habits, metabolic profiles, and environmental adaptability in *Rosa*. For instance, the iconic double flower phenotype, characterized by more than ten petals, emerged convergently in Asia and Europe through selection on the restricted expression of the rose ortholog of AGAMOUS (Dubois et al., 2010). Deciphering the genetic basis underlying this diversity is thus crucial for both evolutionary studies and the advancement of rose breeding.

Genomic studies of *Rosa* have revealed high levels of genetic diversity (Raymond et al., 2018; Hibrand Saint-Oyant et al., 2018; Smulders et al., 2019). Yet, conventional comparative genomics, which typically relies on alignment to a single reference, does not capture the full spectrum of species-wide variation and is prone to reference bias. In contrast, current pangenomic frameworks integrate multiple assemblies into graph-based models, enabling comprehensive identification and characterization of the extensive genetic diversity. This approach is particularly advantageous for dissecting the genetic determinants of complex traits and for uncovering valuable alleles in genetically diverse taxonomic groups such as the genus *Rosa*. The latter has been demonstrated by studies in other Rosaceae species, where the pangenomic approach has unraveled the genetic basis of fruit color and domestication history for strawberry (Qiao et al., 2021) and apple (Sun et al., 2020b).

Recent advances in rose genomics have resulted in several *Rosa* genomes from both wild and cultivated species (Raymond et al., 2018; Hibrand Saint-Oyant et al., 2018; Chen et al., 2021; Zong et al., 2024; Zhong et al., 2021; Zhang et al., 2024; Shang et al., 2024), offering valuable insights into the genetic architecture and evolution of key traits. For example, the evolution of recurrent blooming in cultivated roses has been revealed with the help of the first rose genomes (Raymond et al., 2018; Hibrand Saint-Oyant et al., 2018). While these resources provide the foundation for constructing a comprehensive *Rosa* pangenome, they are predominantly derived from major cultivated lineages and their closely related wild progenitors. Many wild species, especially those representing the basal *Rosa* clade (including sections Hulthemia, Hesperhodos, and Pimpinellifoliae) (Debray et al., 2022) are not represented, even though genomes from these basal species hold the potential to reveal deeper evolutionary insights and novel genetic variations (Khan et al., 2020; Cheng et al., 2025).

Here we present *de novo* genomes for *R. sericea* (section Pimpinellifoliae) and *R. rugosa* based on PacBio HiFi sequencing and Bionano optical mapping. By integrating these high-quality genomes with available genomic resources, we constructed a Rosaceae panproteome and a *Rosa* pangenome (the latter spanning wild, traditional garden, and modern rose lineages) using a De Bruijn graph (DBG) approach. This work addresses the following objectives: (1) characterizing gene family evolution, including contraction and expansion, across the Rosaceae family, with particular emphasis on the *Rosa* lineage, using panproteome analysis; (2) characterizing the genetic variations within the *Rosa* pangenome, with emphasis on gene content dynamics and transposable element (TE)-mediated genomic innovation; (3) reconstructing the evolutionary history of the *NUDX1* gene family, critical for scent biosynthesis, to evaluate the interplay between TE activity and lineage-specific gene family expansions; and (4) addressing technical challenges in pangenome analyses, such as distinguishing biological variation from technical variation, assembly/annotation artifacts, and achieving high-quality genomes through a pangenomic approach. This study not only advances our understanding of *Rosa* genome evolution and adaptive processes but also provides a valuable genomic framework for future research and breeding efforts.

## Results

### Two *de novo* genomes

To expand the genomic resources for the *Rosa* genus, we first generated reference genomes for two distinct diploid roses: *R. rugosa* and *R. sericea. R. sericea* was selected as it occupies a basal position within the *Rosa* genus, for which the genomic resources are currently limited (Wissemann et al., 2003; Wissemann and Ritz, 2005; Fougère-Danezan et al., 2015). *R. rugosa* was selected as an important progenitor species and a source of resistance genes for modern roses (*Rosa hybrida*). *K*-mer analysis confirmed that both species are diploid (Fig. S1). Using PacBio HiFi sequencing (Table S1), Bionano optical mapping, and reference-guided scaffolding, we assembled chromosome-scale genomes for both species (Fig. 1, Table S2). For *R. rugosa*, the high coverage of HiFi data enabled a phased assembly, resolving two haplotypes (hapA: 440 Mb; hapB: 435 Mb). The unphased *R. sericea* genome assembly spanned 356 Mb. These sizes closely matched the estimates obtained from both *k*-mer and flow cytometry analyses (Fig. S2).

**Figure 1:**
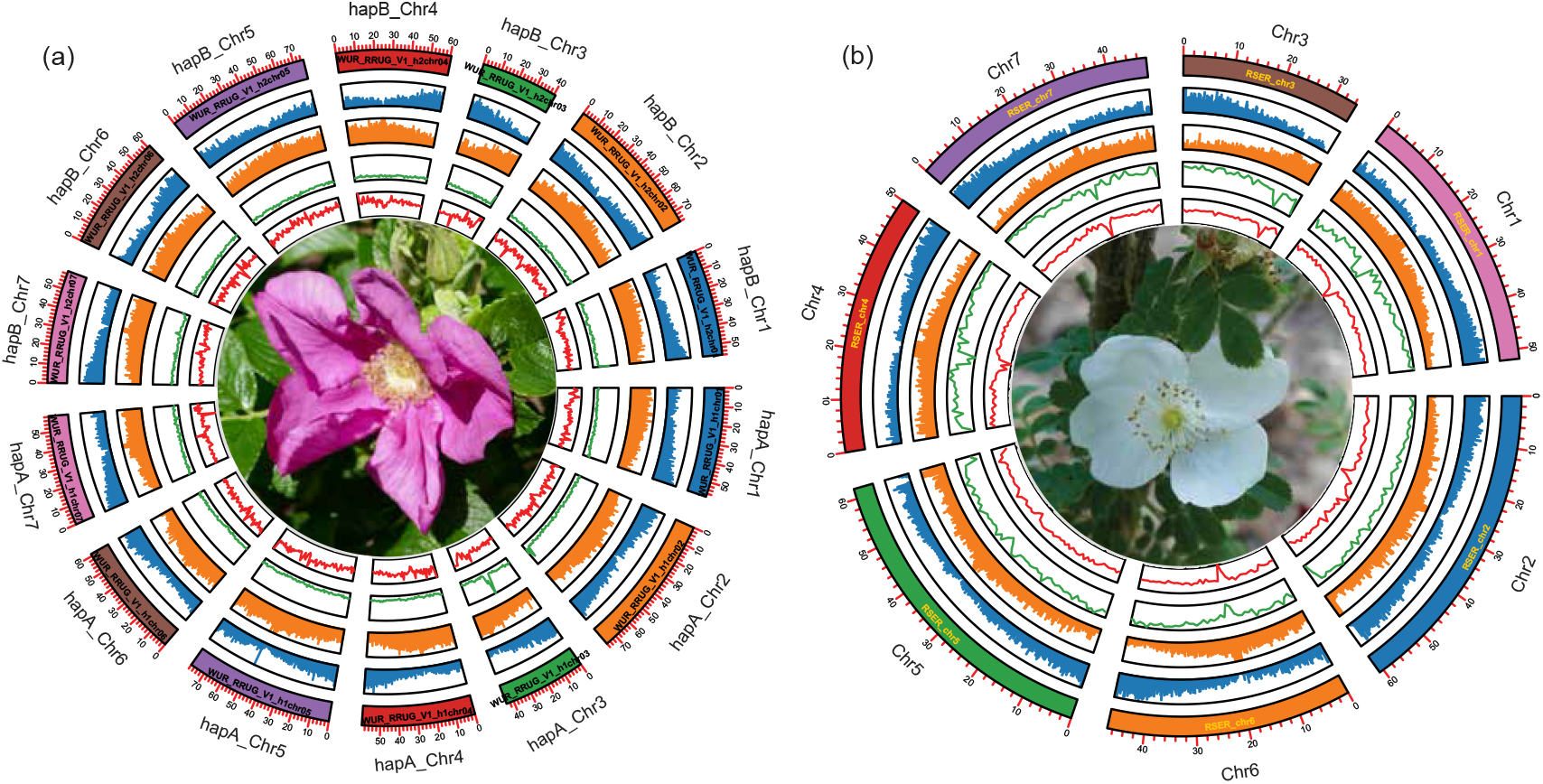
Circos plots for *R. rugosa* and *R. sericea* genomes. (a) Phased *R. rugosa* genome. (b) *R. sericea* genome. Tracks from inner to outer: GC content, Coverage depth of HiFi sequencing reads, TE abundance, Gene content, Chromosomes. Unanchored sequences are not shown.

The genomes of *R. rugosa* and *R. sericea* demonstrated robust quality across multiple metrics (Table S2). High HiFi read and Iso-Seq transcript mapping rates (*>*98% for both species) and near-complete Benchmarking Universal Single-Copy Orthologs (BUSCO) scores (*>*97.7%) confirmed assembly accuracy and completeness. Notably, low BUSCO duplication rates (≤4.3%) and long terminal repeat (LTR) Assembly Index (LAI) values surpassing the gold-quality threshold indicated minimal redundancy and high contiguity (Ou et al., 2018). By searching for the canonical plant telomeric repeat sequence TTTAGGG, we identified all the telomeres in the hapA and hapB of *R. rugosa* genomes (Fig. S3). For *R. sericea*, however, chromosome3 (Chr3) and Chr4 each lack one telomeric region. Merqury analyses confirmed high base-level accuracy, with consensus quality (QV) of 59.84, 61.12, and 59.34 (Table S2). Additionally, a *k*-mer spectrum analysis was used to assess the phasing quality for *R. rugosa* genome (Fig. S4). It was observed that most *k*-mers in the heterozygous peak appeared only once, while those in the homozygous peak were present twice, suggesting little assembly redundancy. Furthermore, we identified balanced haplotype-specific *k*-mers from hapA and hapB, indicating robust phasing quality. These results collectively demonstrate the high accuracy and completeness of the genome assemblies for *R. rugosa* and *R. sericea*.

The genome of *R. sericea* comprises 174.05 Mb of repeat sequences, representing 48.9% of its total genome size, whereas *R. rugosa* exhibits a higher repeat content, with 250.59 Mb (56.9%) in hapA and 247.34 Mb (56.8%) in hapB. This difference aligns with the observed genome size variation between the two species. In both species, LTRs are the predominant repeat element, constituting between 28.5% and 39.2% of the genome (Table S2). Gene prediction identified 30,826 and 30,512 protein-coding genes in the hapA and hapB genomes of *R. rugosa*, respectively, while *R. sericea* was predicted to contain 27,674 protein-coding genes. These gene counts per haploid genome are in line with the RefSeq annotations available for other *Rosa* species (O’Leary et al., 2016). BUSCO analyses of the annotated gene sets revealed high completeness, ranging from 98.1% to 99.1%, while 89.2% and 91.8% of the genes from *R. rugosa* and *R. sericea* could be functionally annotated (Table S2). These results underscore the high quality of gene prediction for both genomes.

### Rosaceae panproteome

To establish the phylogenetic relationships of *Rosa* species within the genus as well as in the broader context of the Rosaceae family, we performed a panproteome analysis using the protein sequences from 14 genome assemblies of Rosaceae species. The dataset included seven *Rosa* species (one representative per species) and six other Rosaceae species. *Ziziphus jujuba* from the Rhamnaceae family was included as an outgroup (Table S3). To standardize the analyses, we retained proteins from the haplotype with the highest BUSCO completeness for phased genomes. However, due to its hybrid nature and the likely abnormal meiosis—evidenced by its pollen abortion phenotype—we included both haplotypes of *R. sterilis* to account for its genome complexity (Zong et al., 2024).

A total of 513,063 protein sequences were clustered into 88,984 homology groups, with 7,640, 27,609, and 53,735 homology groups identified as core, accessory, and unique groups, respectively (Fig. 2). The accessory groups predominantly demonstrated clade-specific patterns; however, some inconsistencies were noted. For example, accessory groups lacking only in *R. sterilis* hapB were more abundant than those missing in *Z. jujuba*, suggesting lower annotation or assembly quality for *R. sterilis* hapB. Unique homology groups can signify biologically meaningful genes but frequently result from annotation artifacts. For instance, *F. vesca* exhibited the highest number of unique homology groups, surpassing even the outgroup species *Z. jujuba*. Within *Rosa, R. hybrida* and *R. sterilis* hapA had over twice the number of unique homology groups compared to other *Rosa* genomes. Finally, the number of core genes in the 7,640 core groups is stable across species, except for two Maleae species, whose variability likely reflects a lineage-specific whole-genome duplication (WGD) event (Fig. 3b).

**Figure 2:**
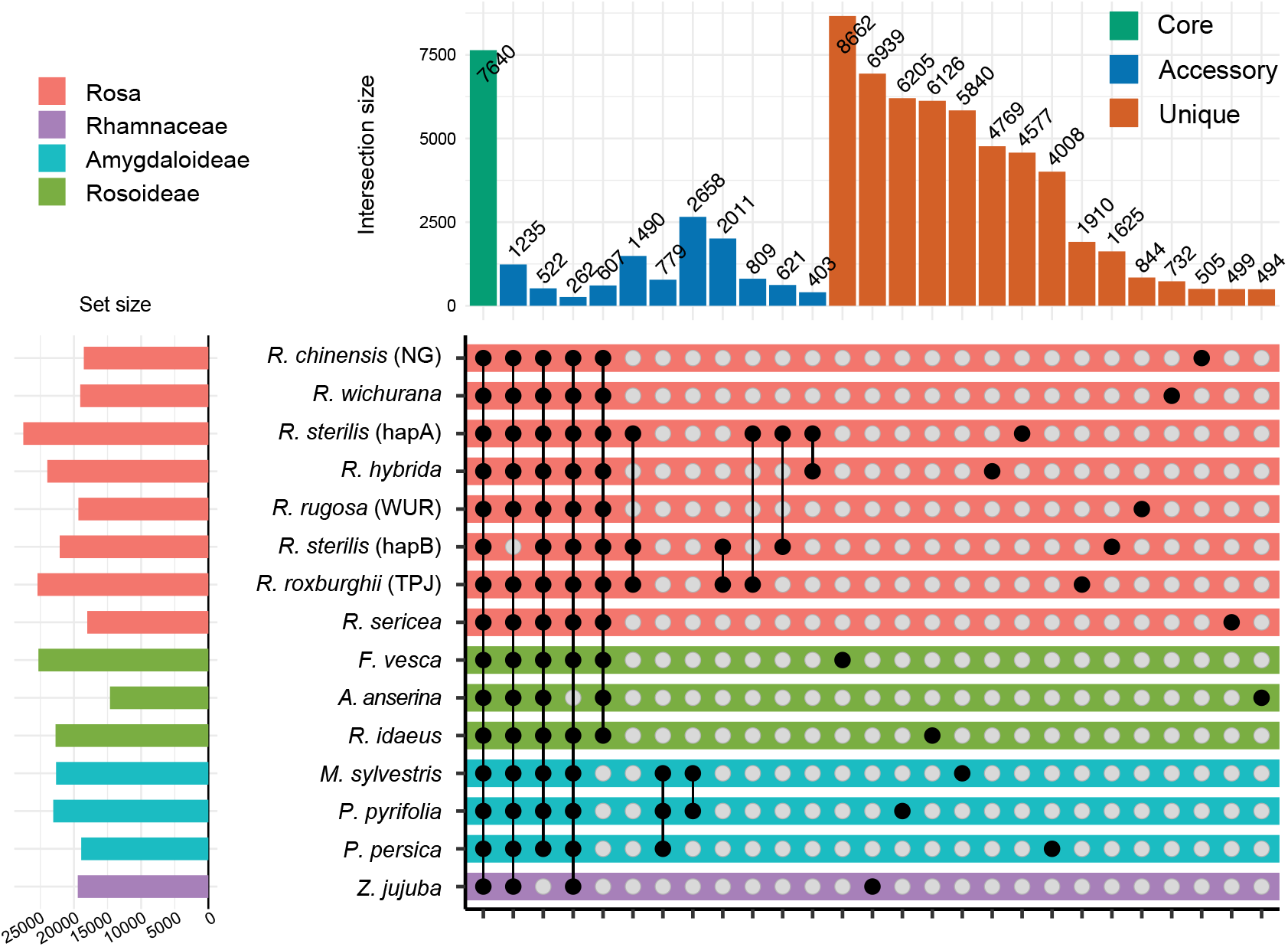
Upset plot illustrating the distribution of core, accessory, and unique homology groups across 14 Rosaceae species. The intersections were sorted by degree for the three types of homology groups. Genomes were ordered by taxonomy. Intersections with size <250 were not shown.

**Figure 3:**
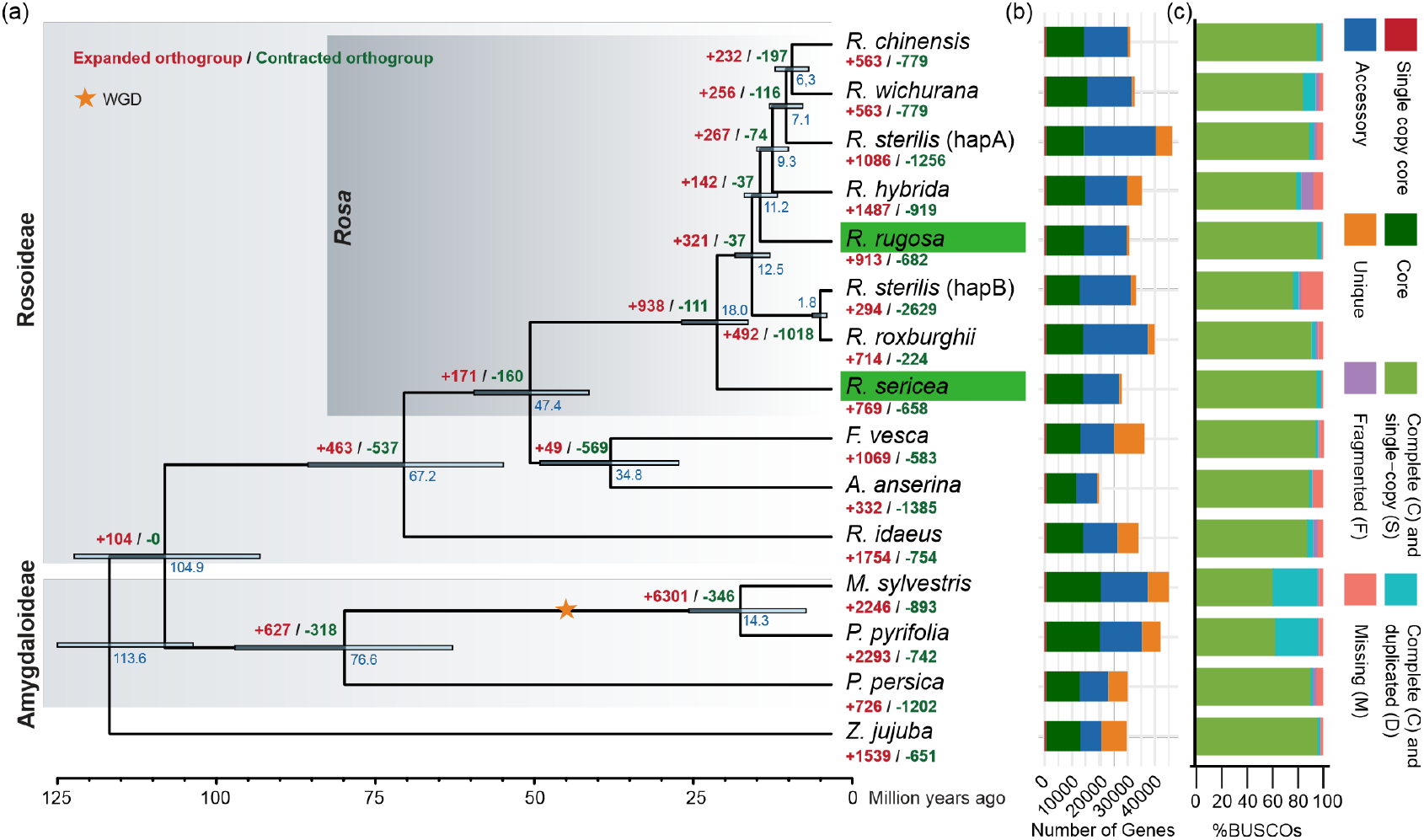
Phylogenomics of Rosaceae species. (a) Phylogenetic analyses of sampled Rosaceae genomes. The two *de novo* genomes reported in this study are indicated by a green background. The numbers of expanded and contracted homology groups are indicated with + and − signs. Numbers at branches (in blue) indicate the estimated mean divergence times (in M years), and grey bars indicate 95% HPD intervals. (b) Gene classification of proteomes. (c) BUSCO completeness of the input proteomes.

We explored the organismal phylogeny by constructing a protein-aligned nucleotide-based species tree based on the full spectrum of core homology groups. This phylogeny robustly divided Rosaceae species into two major subfamilies — Rosoideae and Amygdaloideae — consistent with previous studies that were based on fewer sequences but a broader range of taxa (Xiang et al., 2016; Zhang et al., 2017). Within *Rosa, R. sericea* is resolved as sister to all other sampled *Rosa* species (Fig. 3a). Notably, the two haplotypes of *R. sterilis* were placed distantly: hapB clustered with the two *R. roxburghii* genomes while hapA formed a sister group to cultivated roses, confirming the hybrid origin of *R. sterilis* (Zong et al., 2024). Through molecular dating constrained by fossil evidence, we estimated the divergence time for the constructed Rosaceae phylogeny. The crown age for Rosaceae was estimated at approximately 104.9 million years ago (Mya). Within Rosoideae, the *Rosa* lineage diverged from other members at approximately 47.4 Mya, and the crown age of the sampled *Rosa* species was estimated to be about 18.0 Mya, indicating their recent diversification during the Miocene (Fig. 3a). These results agree with previous studies (Xiang et al., 2016; Su et al., 2016; Zhang et al., 2017; Debray et al., 2022).

We next traced the contraction and expansion of homology groups across the constructed Rosaceae phylogeny. At the species level, the two Maleae species, *Malus sylvestris* and *Pyrus pyrifolia*, exhibited the highest number of expanded homology groups, likely reflecting the impact of the Maleae-specific WGD event (Velasco et al., 2010; Wu et al., 2013) (Fig. 3a). Within the genus *Rosa*, the hapB of *R. sterilis* showed the highest number of contracted homology groups (2,629), which is likely an artifact due to its incomplete proteome, as reflected by its low BUSCO completeness (Fig. 3a,c). Functional enrichment suggested that most of the expanded and contracted homology groups at species-level in *Rosa* are linked to secondary metabolism, response to environmental factors, and DNA/RNA replication (Fig. S5, S6) (Table S4, S5). Interestingly, we observed clade-specific differences in the functional enrichment of the expanded homology groups. In Amygdaloideae, the enriched groups were predominantly associated with transporter activity and transcription regulation, whereas in *Rosa* they were predominantly linked to secondary metabolism, particularly in the biosynthesis of monoterpenoids, unsaturated fatty acids, sesquiterpenoids, and steroids. These compounds are believed to play roles in plant adaptation and contribute to the characteristic scent of *Rosa* species (Conart et al., 2023; Zhou et al., 2024) (Fig. S7, S8) (Table S6, S7).

### *Rosa* pangenome

To construct a *Rosa* pangenome, we integrated our two *de novo* genomes with ten additional publicly available chromosome-level, high-quality genomes, forming a representative dataset spanning wild, old garden, and modern rose lineages (Table S3). Pairwise mash distance analysis revealed substantial nucleotide-level divergence across the genus, with the highest genetic distance (0.06) observed between *R. sericea* and *R. roxburghii* (BMC), indicating extensive sequence diversity within *Rosa* and confirming the basal position of *R. sericea* within the genus (Fig. S9). Alignment-based pangenome methods often struggle with extensive sequence variation, leading to alignment inaccuracies or un-alignable regions. PanTools (Jonkheer et al., 2022), a generalized DBG-based pangenomic tool, circumvents this issue using *k*-mers, and was therefore selected to construct the *Rosa* pangenome.

Classification of the *Rosa* pangenome revealed a greater number of core homology groups compared to the broader Rosaceae panproteome, which is expected given the lower taxonomic level. Applying Heap’s law yielded an estimated alpha of 0.30, indicating an open pangenome where novel genes continue to emerge as additional genomes are sampled (Tettelin et al., 2008). It is important to note that the total number of homology groups may be somewhat inflated due to potential false gene predictions. An examination of the intersection of accessory homology groups revealed substantial gene conservation between the *R. roxburghii, R. sterilis*, and *R. hybrida* genomes, mirroring patterns observed in the broader Rosaceae panproteome analyses (Fig. S10).

Compared to traditional phylogenetic methods that rely on a limited number of genes, our pangenomic approach leverages 4,367 single-copy core homology groups that appeared in all haplotypes, providing a comprehensive framework for robust evolutionary inference (Fig. 4). The 18 *Rosa* haplotypes clustered based on their domestication status, except for the hapA of *R. sterilis*, which is derived from the *Rosa longicuspis* lineage (Zong et al., 2024). Notably, the topology of the haplotype-level *Rosa* phylogeny differed slightly from that of the Rosaceae panproteome-derived tree (Fig. 3, 4). This difference likely arises from hybridization-driven genome mosaicism in cultivated roses as multiple *R. chinensis* haplotypes were included. Additionally, all four haplotypes of *R. hybrida* form a sister group to the old garden roses, suggesting its genetic architecture is largely derived from these traditional cultivars (Zhang et al., 2024). In particular, the phased *R. chinensis* (MH) haplotypes diverge: hapA clustered with other *R. chinensis* genomes, while hapB grouped with *R. wichurana*. This clear split further corroborates the extensive hybridization that has shaped the genome of cultivated roses (Raymond et al., 2018; Zhang et al., 2024).

**Figure 4:**
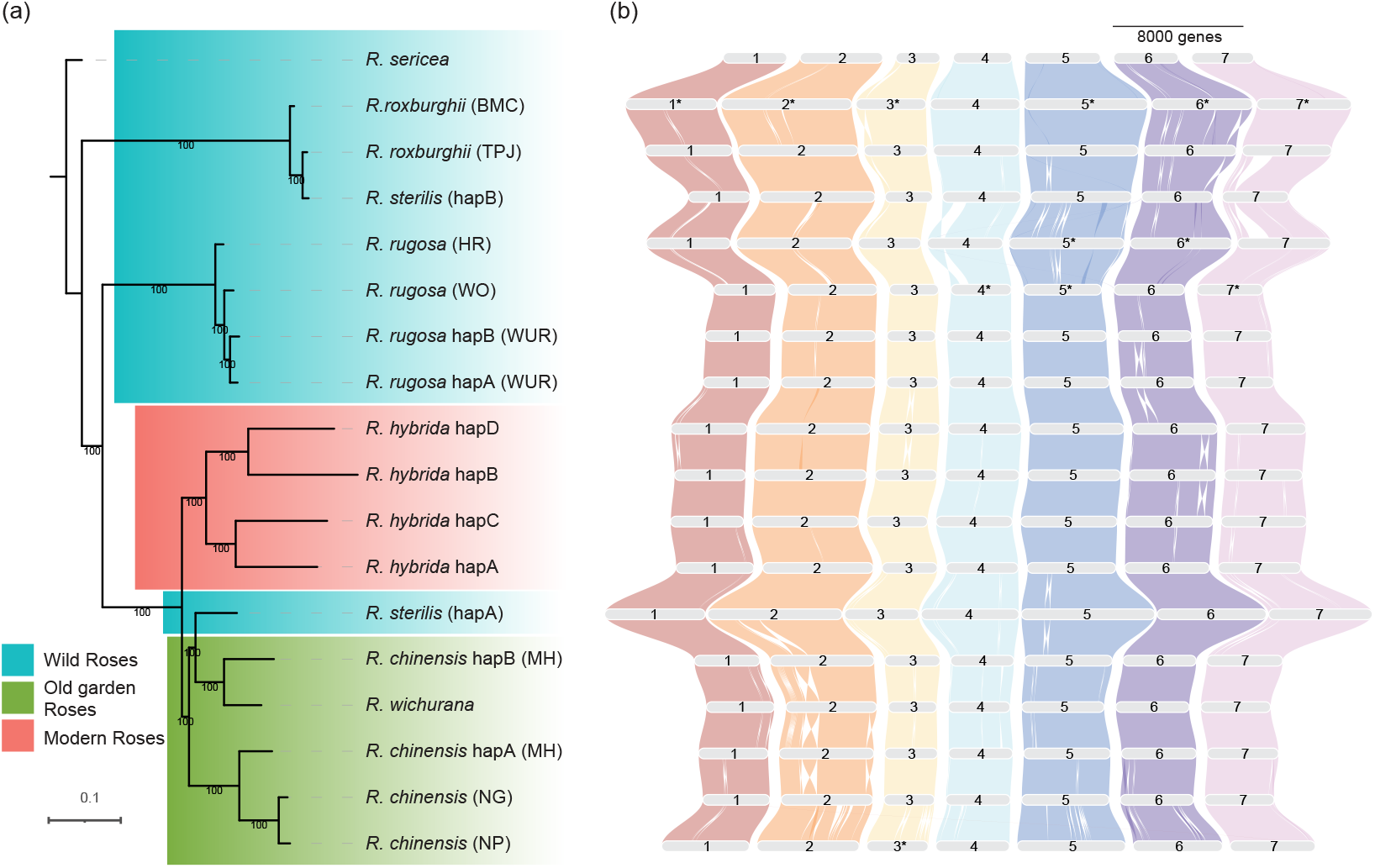
Phylogeny of the *Rosa* pangenome. (a) Maximum likelihood phylogeny of haploid *Rosa* genomes based on SNPs identified in single-copy orthologous genes. Node numbers indicate % bootstrap support (1000 replicates). (b) Gene-based macrosynteny across the pangenome. The *R. chinensis* (NG) genome was used as reference for coloring and ordering of chromosomes. Scale bar: 8,000 genes. Chromosomes for which the orientation conflicted with the consensus orientation were re-oriented and denoted with an asterisk (*).

Gene-based synteny (Fig. 4b), as well as alignment-based synteny (Fig. S11), showed that the chromosome structure is conserved across the *Rosa* genus, as no obvious interchromosomal rearrangements were observed. However, several structural variants exist. For example, Chr2 of both haplotypes of *R. chinensis* (MH) share a unique large inversion and Chr6 of hapA from *R. rugosa* (WUR) exhibits a haplotype-specific inversion. Some structural variants are likely assembly artifacts, such as the missing part of Chr1 in hapB of *R. sterilis* and the missing part of Chr2 in *R. rugosa* (HR) (Fig. S11). These assembly issues contributed to reduced genome completeness, reflected by lower BUSCO scores at the haplotype level (Fig. S12).

To accommodate multiple genomes per species and fully leverage the available data, we redefined core, accessory, and unique homology groups at the species level rather than the genome level. Core gene models, which are supported by multiple independent sequencing efforts, are considered the most reliable, while unique genes — although potentially revealing novel biology — are expected to contain a higher rate of annotation errors and should be interpreted with caution. Accessory groups, which receive moderate support, might also include some annotation artifacts. By comparing gene metrics across homology group types, it was found that core genes exhibited longer lengths and higher exon numbers compared to accessory and unique genes (Fig. S13b,c). Accessory and unique genes showed high single-exon prevalence, suggesting either annotation issues or recent evolutionary origins (Fig. S13d). In addition, core genes exhibit slightly higher GC content than accessory genes, while unique genes show greater variation in GC content (Fig. S13a).

We next focused our analyses on the core genes, since these are considered more reliable than accessory or unique genes. Copy number variation analysis revealed substantial lineage-specific expansions (Fig. S14). Interestingly, homology groups that are expanded in *R. chinensis, R. hybrida* and *R. rugosa* but not in the more basal species *R. sericea, R. roxburghii* and *R. sterilis* were significantly functionally enriched in pathways including monoterpenoid biosynthesis, lipid metabolism, flavonoid biosynthesis, benzoxazinoid biosynthesis and phenylpropanoids biosynthesis (*p* < 0.05) (Table S8, S9). These pathways are closely linked to rose scent production, cold acclimation, and sugar metabolism.

*R. sericea* also showed some interesting lineage-specific expansion in core gene families (Fig. S14), despite its compact genome (smallest genome size and lowest gene count among the sampled *Rosa* genomes). These species-specific expansions were significantly linked to environmental adaptation and anthocyanin biosynthesis. Notably, gene families such as acylamino-acid-releasing enzymes, short-chain dehydrogenase/reductases, and F-box proteins—key players in oxidative stress response— were expanded (Table S10, S11). Moreover, *R. sericea* exhibited an expansion of UGT88a1 genes compared to the other genomes. Given that UGT88a1 plays a crucial role in the glycosylation of flavonoids like quercetin (Lim et al., 2004), a key mechanism for mitigating stresses such as UV exposure, this expansion likely contributes to enhanced stress tolerance. Together, these findings reveal a potential genetic mechanism underlying the environmental adaptation of *R. sericea* to high altitudes (up to 4,000 m) (Gao et al., 2019).

### Transposable element evolution across *Rosa*

Transposable elements (TEs) are key drivers of plant genome evolution, directly influencing genome size variation and modulating gene regulatory networks that shape phenotypic diversity (Pulido and Casacuberta, 2023; Bennetzen and Wang, 2014; Yu et al., 2024). In the *Rosa* pangenome, we investigated the contribution of TEs to genome size variation and evolution. Genome size variation of haploid *Rosa* genomes correlates with TE content. For instance, *R. sericea* (356 Mb genome) has the lowest TE abundance (210 Mb) while the largest genome *R. chinensis* (MH) hapB (541 Mb) contains the highest TE content (358 Mb). This aligns with typical genomic patterns, where TE expansion drives genome size differences (Fig. 5a) (Table S12). Among the major TE classes, PIF/Harbinger, Gypsy, Copia, and CACTA showed strong correlations with genome size (Fig. S15).

**Figure 5:**
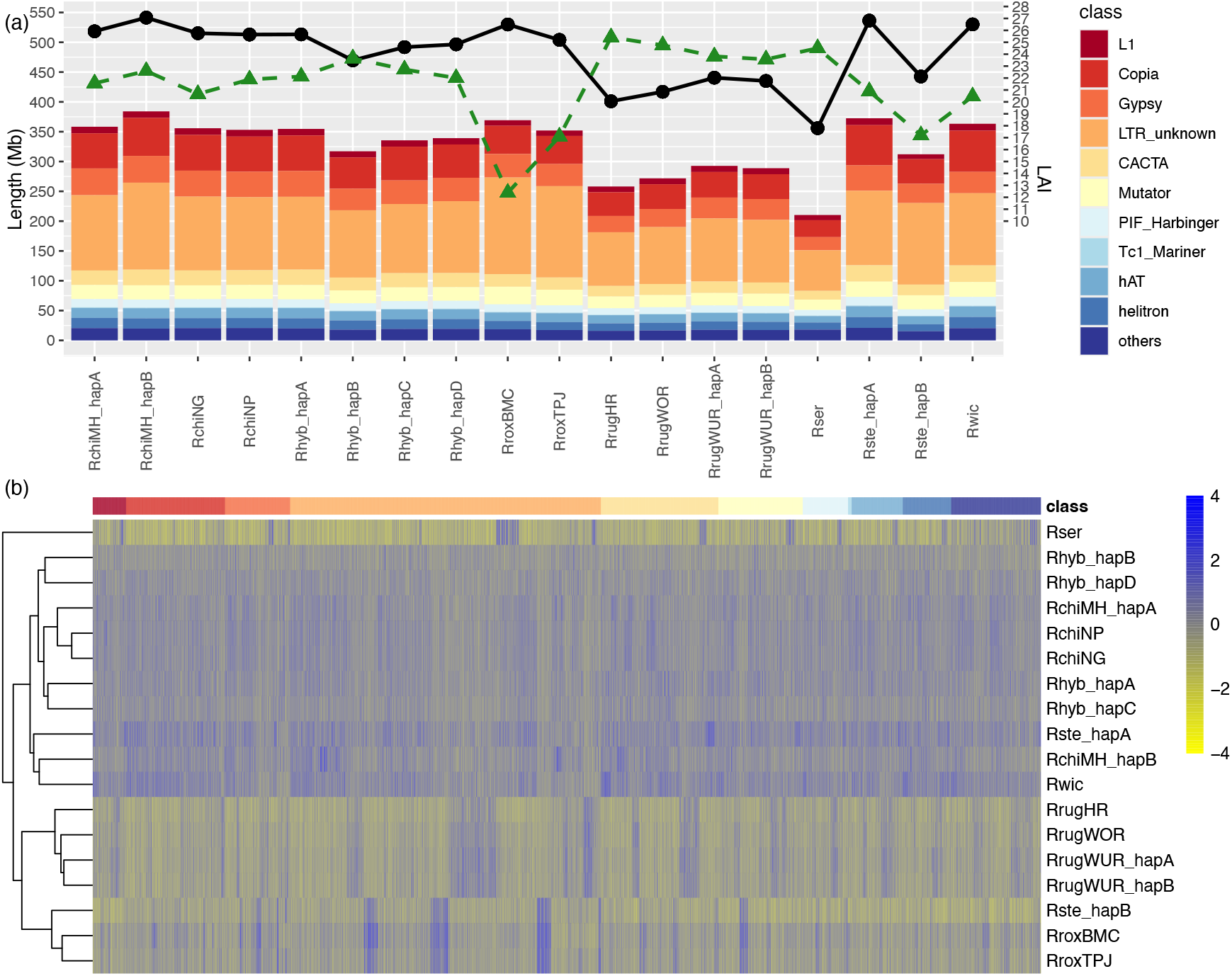
Transposable Element (TE) abundance in the *Rosa* pangenome. (a) TE types and content of *Rosa* haploid genomes. The black dots and solid line indicate genome sizes for each haplotype assembly. The green triangles and dashed line indicate LTR Assembly Index (LAI) values per each haplotype assembly. (b) Abundance heatmap of TE families across the *Rosa* pangenome. The top annotation bar shows each family’s class. For each TE family, member counts were standardized using Z-scores to facilitate direct comparison of their relative abundance.

The abundance of TEs exhibited a pronounced species specificity in wild roses: *R. sericea* showed depletion across nearly all major TE classes, whereas *R. rugosa* and *R. roxburghii* displayed lineage-specific expansions mainly in LTR, CACTA, and Mutator TEs (Fig. 5b). In contrast, cultivated roses (old and modern) shared remarkably similar TE profiles, suggesting no marked TE bursts or suppression during domestication. This pattern implies that human selection over the past thousands of years has not yet substantially reshaped TE landscapes, likely due to the short timescale of domestication relative to TE evolutionary dynamics (Bendahmane et al., 2013). Repeat fingerprints differ between homologous haplotypes, mirroring their phylogenetic origins. For example, the hapB of *R. sterilis* shares a long terminal repeat (LTR unknown) expansion pattern with *R. roxburghii* genomes, consistent with their close evolutionary relationship.

We then focused on full-length LTRs, as these provide evidence for evolutionary dating. Most *Rosa* genomes exhibit a very recent burst of LTR activity (identity *>* 0.995) estimated at around 0.63 Mya (using the Rosaceae mutation rate estimate of 4 × 10^−9^ mutations per site per year (Sun et al., 2020b)). On the contrary, the two *R. roxburghii* genomes and hapB of *R. sterilis* showed a different pattern, with LTRs of unknown identity centered around 0.985 to 0.990 (roughly 1.26-1.89 Mya) (Fig. S16). These findings indicate that the LTR burst likely occurred in the common ancestor of the *R. roxburghii* lineage, before the hybridization event that led to the emergence of *R. sterilis* (Zong et al., 2024).

Chromosomal organization suggests that TEs are distributed non-randomly, with their abundance dropping sharply from pericentromeric heterochromatin to the euchromatic arms, reflecting the inverse relationship between TE density and gene density (Fig. S17, S18). The conserved TE distribution pattern across the *Rosa* pangenome also enables us to identify the putative centromeres. Notably, Gypsy-type LTRs dominate centromeric regions, consistent with their role in maintaining centromere identity in plants (Fig. S17, S18) (Neumann et al., 2021). This TE compartmentalization mirrors patterns in other Rosaceae genomes including *F. vesca* and *M. domestica* (Jin et al., 2025; Su et al., 2024), suggesting deep conservation of TE-chromatin interactions across Rosaceae.

### Pangenomic analysis of NUDX1 evolution

Among the gene families exhibiting lineage-specific copy number variations, the NUDX1 family stands out because of its suggested role in geraniol formation — the major compound responsible for rose scent. Previous studies have proposed that the *NUDX1* family emerged in the common ancestor of the Rosoideae and subsequently evolved via multiple tandem duplications (Magnard et al., 2015; Sun et al., 2020a; Conart et al., 2022). Phylogenetic analysis revealed that NUDX1 cluster into three well-separated groups: NUDX1-3, NUDX1-2, and NUDX1-1, with NUDX1-2 and NUDX1-1 further subdivided into three and two subgroups, respectively (Fig. 6) (Table S13). While most subgroups are broadly conserved across the *Rosa* pangenome, NUDX1-1a is restricted to *R. sterilis, R. rugosa*, and *R. chinensis*, indicating lineage-specific expansion. Notably, two *R. chinensis* members clustered with *R. rugosa* sequences with NUDX1-1a (with high support; Fig. S19), indicating a shared evolutionary origin.

**Figure 6:**
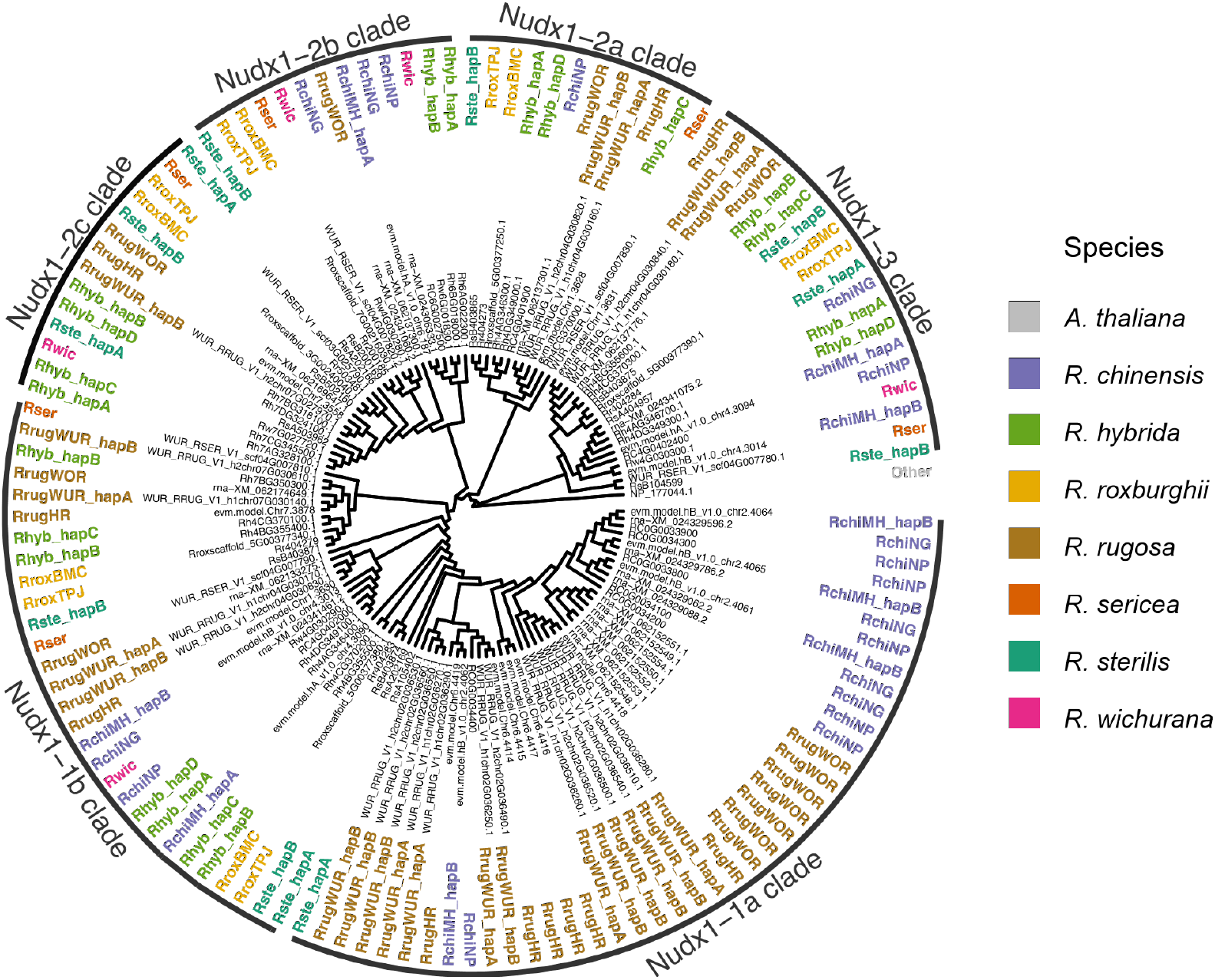
Phylogenetic analysis of the *NUDX1* gene family in the *Rosa* pangenome.

We next investigated the genomic distribution of *NUDX1* members across the *Rosa* pangenome. Our analysis indicates that these genes are located on four chromosomes: Chr2, Chr4, Chr6, and Chr7 (Fig. S20, S21, S22, S23). Notably, Chr4 of *R. sericea* retains an ancestral *NUDX1* cluster (containing an array of *NUDX1-3, NUDX1-2a, NUDX1-2b*, and *NUDX1-1b* genes), suggesting tandem duplications predating *Rosa* lineage divergence. *NUDX1-2c* resides in a highly conserved region on Chr7 across most genomes, where we identified LINE-type retrotransposon fragments from the same family upstream of all *NUDX1-2c* loci. Members from the above LINE family were also present in the tandem *NUDX1* region on Chr4 in early-diverging species (*R. sericea* and *R. roxburghii*) (Fig. S22), suggesting a shared ancestral origin. Despite their shared lineage, the above Chr7- and Chr4-derived LINE fragments contain two distinct putative regulatory motifs: Chr7-derived fragments harbor C2H2 zinc finger factors binding sites, while Chr4-derived fragments feature tryptophan cluster factor motifs (Fig. S24, S25). This divergence implies that lineage-specific LINE element insertions have introduced regulatory innovations that modulate *NUDX1-2c* expression. We propose that *NUDX12c* originated on Chr4 and was transposed to Chr7 via an ancient LINE-mediated event likely predating the *Rosa* diversification, with subsequent regulatory motif differentiation. A similar mechanism may have facilitated the transposition of *NUDX1-1a* from Chr4 to Chr2 via a Copia-type TE and evolving of its tissue-specific expression pattern (Conart et al., 2022).

Despite widespread conservation, *NUDX1-2c* is absent from all *R. chinensis* genomes in our pangenome. However, an alternative assembly of *R. chinensis* that retained heterozygous regions revealed *NUDX1-2c* on one contig, suggesting that it was ancestrally present and subsequently lost in a haplotype-specific manner (Sun et al., 2020a). Similarly, *NUDX1-1a* exhibits restricted distribution: it is retained on Chr4 in *R. rugosa, R. chinensis*, and haplotype A of *R. sterilis*, but absent in *R. wichurana* and *R. chinensis* (MH) hapB. This may be due to hybridization-associated haplotype erosion, or because it was already polymorphic in the ancestral haplotypes. However, incomplete genome assembly cannot be ruled out in both cases. Notably, *R. sterilis* hapA carries a truncated *NUDX1-1a* allele (RsA105402) with abnormal protein length (Fig. S26), likely resulting from misannotation or gene truncation.

All four *R. hybrida* haplotypes lack the *NUDX1-1a* gene. When comparing these regions to those in *R. rugosa* and *R. chinensis*, which do carry *NUDX1-1a*, we observed notable dissimilarities (Fig. S23). Notably, hapB and hapD of *R. hybrida* exhibit high genomic similarity and share conserved LTR insertions with *R. wichurana*, a species that also lacks *NUDX1-1a* (Fig. S23). This distinct genomic architecture implies that *R. hybrida* inherited genomic regions devoid of *NUDX1-1a* from ancestral progenitors, rather than having lost the gene due to domestication-associated selection or genomic instability.

## Discussion

### Building a *Rosa* pangenome

In this study, we present two *de novo Rosa* genomes along with a comprehensive *Rosa* pangenome that highlights extensive genomic diversity within the genus, including substantial variations in gene content, structural rearrangements, and TE dynamics. Notably, we demonstrate that the DBG-based pangenomic approach is an efficient approach for capturing and characterizing genomic variation across species with large genetic distances. Alignment-based graph methods such as PGGB and Minigraph-Cactus (Garrison et al., 2024; Hickey et al., 2024) rely on sequence homology to identify syntenic regions, a strategy that fails when genetic distances exceed alignment thresholds or when structural variation disrupts genomic collinearity - common challenges in the highly heterozygous and diverse *Rosa* genomes.

Although gene-based pangenome approaches have been successfully applied to other genera, including *Malus* and *Populus* (Wang et al., 2023; Shi et al., 2024), these approaches primarily focus on the gene layer. Consequently, they capture only part of the genomic variation and cannot easily map those variants back onto the underlying genomic sequences. By constructing the pangenome graph directly from *k*-mers, our approach overcomes these limitations by capturing the complete spectrum of variation present in the sampled genomes. Additionally, the incorporation of annotation-derived homology information enhances our ability to robustly detect and distinguish both conserved and lineage-specific genomic features, providing deeper insights into the evolutionary dynamics shaping the *Rosa* genus.

### Quality discrepancies highlighted by pangenome

Our analyses of the Rosaceae panproteome and the *Rosa* pangenome revealed notable quality discrepancies likely originating from inconsistencies in genome assembly and gene annotation across studies. Specifically, the genomes of *R. roxburghii* (BMC) and *R. hybrida* show an unusually high number of unique homology groups, likely reflecting potential inflation driven by redundant gene models and fragmented contigs, rather than genuine biological novelty.

Furthermore, the homology group classification suggested that *R. hybrida* and *R. roxburghii* genomes share a notably high number of exclusive homology groups. While this pattern could indicate shared evolutionary pressures or similar functional constraints, in this case it does not reflect direct phylogenetic relatedness since *R. roxburghii* is not considered a direct progenitor of *R. hybrida* (Zhang et al., 2024). Supporting evidence from gene-based phylogeny and TE dynamics further substantiates this interpretation. As an alternative, the observed similarity may be the result of systematic annotation biases shared between these genomes.

Substantial differences between the two haplotypes of *R. sterilis* were observed. Specifically, hapA and hapB displayed notable disparities in the panproteome upset analysis, and synteny-based comparisons suggest several segmental deletions in hapB. These observations may reflect genuine biological variation, such as heterozygous structural deletions, but could equally indicate assembly errors or annotation inaccuracies, or possibly a combination thereof.

Clearly distinguishing technical artifacts from true biological variation is essential for accurate interpretation. Our pangenomic approach effectively identifies genuine biological diversity while simultaneously highlighting technical inconsistencies. To address these challenges, we propose a standardized quality control framework for future *Rosa* genome projects. This framework should include consistent naming conventions, mandatory provision of raw data for independent quality assessments, and alignment of genome annotations with established gold-standard gene predictions, such as those available for *R. chinensis* (NG) and *R. rugosa* (WUR). Such standardized practices will significantly enhance the reliability and interpretability of comparative genomic analyses in *Rosa*.

Despite these technical challenges, our results demonstrate that the pangenome provides a robust and informative framework for comparative genomics research in *Rosa*.

### Phylogenetic Insights

Our panproteome analyses estimated the crown age of the sampled *Rosa* species at approximately 18.0 Mya, with *R. sericea* as a basal lineage. This Miocene origin is consistent with analyses based on nuclear single-copy orthologous tags, which dated the *Rosa* crown to around 21.5 Mya (Debray et al., 2022), although previous studies using chloroplast markers have yielded slightly older estimates (~30 Mya) (Fougère-Danezan et al., 2015). The differences among these estimates likely reflect the distinct evolutionary rates and inheritance patterns of nuclear versus chloroplast genomes.

At the species level, the phylogenetic trees reconstructed from both panproteome and pangenome data produced largely congruent topologies. However, discrepancies emerged when considering haplotype-level relationships; for example, the two haplotypes of *R. chinensis* (MH) were placed in separate groups. This pattern likely reflects the extensive hybridization and recombination during the evolution and domestication of cultivated roses, resulting in mosaic genomes that retain localized polymorphisms. Such mosaicism has been reported in previous studies and may be central to understanding the evolution of complex traits in *Rosa* (Raymond et al., 2018; Hibrand Saint-Oyant et al., 2018; Smulders et al., 2019; Zhang et al., 2024). Future efforts on integrating additional sequencing data from putative progenitor species would further elucidate these patterns and their impact on trait diversification.

### Evolution dynamics of gene content

One of the key findings of our study is the substantial copy-number variation observed in core homology groups across *Rosa* species. By focusing on core genes, which offer a more reliable basis for comparative analyses than dispensable genes, we uncovered clear patterns of gene family expansion and contraction linked to both evolutionary history and species-specific adaptations.

Our analysis demonstrated that the NUDX1 family, which is essential for producing scent-related volatiles, resides within a conserved genomic region. The emergence of *NUDX1-1a* appears to have occurred in the common ancestor of *R. rugosa* and *R. chinensis*, with subsequent expansions in these species arising either independently or through concerted evolution (Liao, 1999). Reticulation and frequent hybridization events, as seen in *R. hybrida*, have maintained localized polymorphisms, potentially explaining the observed variation in geraniol profiles among descendant species.

Moreover, our findings highlight a strong association between TE activity and the diversification of the *NUDX1* family. TEs are thought to exhibit non-random insertion patterns influenced by genomic context and selection pressures (Bourque et al., 2018). Their recurrent enrichment near the *NUDX1* loci suggests that TE-mediated structural variation may have facilitated functional diversification of *NUDX1* family in *Rosa*. Similar mechanisms have been reported in other Rosaceae species, where TEs were involved in regulating allele-specific expression patterns (Tian et al., 2022; Yu et al., 2024). Further functional studies are needed to disentangle whether TE activity at *NUDX1* loci reflects adaptive processes, neutral evolution, or a combination of both.

### *Rosa* genome evolution shaped by TE dynamics

Our analysis underscores the importance of TEs in shaping the evolution of the *Rosa* genomes. We observed that TE content varies considerably among species, contributing to genome size differences. Additionally, the enrichment of Gypsy-type LTR retrotransposons in (peri)centromeric regions of *Rosa* genomes underscores their functional role in centromere evolution, potentially mediated through epigenetic regulation (Neumann et al., 2011). In particular, the bimodal pattern of LTR insertions in *R. roxburghii* suggests that at least two distinct waves of TE activity have occurred. Similar episodic bursts of LTR retrotransposons have been documented in wheat (Li et al., 2022), where these have been linked to environmental stress and shifts in epigenetic regulation. Although we did not directly assess TE removal mechanisms in our study, the diverse LTR insertion profiles observed in the *Rosa* pangenome may indirectly reflect variations in TE silencing responses, potentially driven by environmental factors or changes in genomic regulatory processes.

Interestingly, while wild species exhibited distinct TE abundance patterns compared to cultivated ones, we found no clear difference between old cultivated and modern roses. This suggests that TE accumulation is driven primarily by natural evolutionary processes and environmental adaptation, rather than by domestication. In contrast, in some other crops — such as *Zea mays*, where Gypsy TE content differs substantially between tropical and temperate lines — TE dynamics appear to be more tightly linked to domestication and adaptation over longer time scales (Hufford et al., 2021). These findings suggested that the relatively short domestication process in *Rosa* may not have allowed sufficient time for TE divergence among cultivars.

### Perspectives

Our current *Rosa* pangenome integrates genomic data from both wild and cultivated roses, but primarily represents lineages of Asian origin, leaving European and North American genetic backgrounds underrepresented. While this pangenome effectively captures the core genomic content across *Rosa* species, the Heap’s law alpha value of 0.30 indicates an open pangenome, revealing extensive unexplored diversity within accessory and unique genomic regions. This finding emphasizes the necessity of incorporating additional genomes, particularly from geographically diverse lineages, to fully characterize the genetic breadth of the genus.

Furthermore, observed haplotype diversity, notably the heterozygous retention of *NUDX1-2c*, and the distinct phylogenetic placement of *R. chinensis* (MH) haplotypes highlight the crucial importance of phased genome assemblies. Integrating resequencing data with pangenomic analyses remains an underexploited strategy that could significantly deepen our understanding of genetic diversity. Moving forward, developing expanded pangenomes and complementary multiomics data (e.g. transcriptomics and metabolomics) will be essential for unraveling the evolutionary processes driving trait diversification and adaptation within *Rosa*.

## Conclusions

In summary, we generated two high-quality *Rosa* genomes and integrated them with public resources to construct a *Rosa* pangenome. Our analyses revealed significant genomic diversity among species and highlighted extensive variation in core gene content and TEs that underlie both genome evolution and species-specific adaptations. This is exemplified by our detailed investigation of the *NUDX1* gene family, a key gene in rose scent biosynthesis whose evolution appears to be tightly linked to TE dynamics. Our study demonstrates the advantages of a generalized DBG-based pangenomic approach that integrates genomic sequences and annotations from diverse species. Unlike common alignment-dependent methods which risk losing critical sequence details, our alignment-free strategy preserves full sequence information. This comprehensive framework enables a more accurate exploration of the genetic basis of traits such as scent biosynthesis, stress adaptation, and domestication. Finally, our pangenome not only advances our knowledge of *Rosa* genome evolution but also serves as a valuable resource for future genetic improvement and breeding, with further insights expected from integrating genomic and transcriptomic data from additional roses with diverse genetic backgrounds.

## Methods

### Plant materials and sequencing

Sequencing was carried out on individual plants of *R. rugosa* (accession: 19–BG21966/1) and *R. sericea* (accession: 19– BG24419/1), both cultivated at the Belmonte Arboretum, Wageningen, The Netherlands. High-molecular-weight DNA was isolated from leaf tissues using CTAB method (Porebski et al., 1997). For DNA sequencing, libraries were prepared and sequenced using the PacBio RSII platform in Circular Consensus Sequencing (CCS) mode. Additionally, RNA sequencing (RNA-seq) was conducted on leaf, stem, flower bud, and mature flower tissues, all of which were collected in situ and immediately stored in liquid nitrogen. For Bionano optical mapping, the Bionano Prep Plant Tissue DNA Isolation Kit was used.

### Mitochondria and chloroplast assembly

The complete mitochondrion (GenBank accession: NC_065237.1) and chloroplast (GenBank accession: NC_044094.1) sequences of *R. rugosa* were downloaded from NCBI and used as reference to reconstruct the organellar genomes from the sequenced accessions. HiFi reads were mapped to the reference using minimap2 v2.26-r1175 (Li, 2018) and further extracted for assembly using Samtools v1.18 (Li et al., 2009). To assemble the chloroplast genomes, ptGAUL v1.0.5 (Zhou et al., 2023) was used with default parameters. Script ‘get organelle from assembly.py’ from GetOrganelle v1.7.7.0 (Jin et al., 2020) was used to convert the gfa format to fasta format output. To assemble the mitochondrial genomes, nextDenovo v2.5.2 (Hu et al., 2024b) was used with default parameters. The assembled mitochondrial and chloroplast genomes were polished using organellar HiFi reads using NextPolish2 v0.2.0 (Hu et al., 2024a).

### Genome assembly

Before assembling the nuclear genomes, a strict quality control on HiFi reads was performed. First, the NCBI Foreign Contamination Screen (FCS) v0.5.0, including both the FCS-adaptor and FCS-GX, was used to remove contaminant HiFi reads (Astashyn et al., 2024). Next, reads passing the FCS check were aligned using blast v 2.14.0+ (Camacho et al., 2009) against the assembled mitochondrial and chloroplast genomes, to filter out organellar reads. The following criterion was used to identify organellar HiFi reads: if more than 90 % of a read returned a hit with the reference organellar genomes, it was labeled as an organellar HiFi read. The genome size, heterozygosity, and ploidy level of *R. rugosa* and *R. sericea* were estimated using GenomeScope v2.0 and Smudgeplot v0.2.5 (Ranallo-Benavidez et al., 2020) with nuclear HiFi reads as input.

The nuclear HiFi reads (Table S1) were fed to hifiasm v0.19.5-r587 (Cheng et al., 2022) to perform the assembly in HiFionly mode. Two partially-phased assemblies and primary/alternate assemblies were produced for *R. rugosa* and *R. sericea* genomes. Based on the assemblies’ statistics, the two partially-phased assemblies for *R. rugosa* and primary assemblies for *R. sericea* were selected for further Bionano scaffolding (Lam et al., 2012). For *R. rugosa*, the two haplotypes for each chromosome, produced by hifiasm, were arbitrarily assigned to hapA or hapB, with no inherent biological ordering. For genome scaffolding, optical genome mapping was used. Raw Bionano molecules were first filtered with a minimum length of 150 kb and then assembled *de novo* into optical maps. In parallel, *in silico* maps were generated from HiFi genome assemblies. Based on the optical maps and *in silico* maps, hybrid scaffolding was performed using Bionano Solve v3.8. The contiguity of the hybrid scaffolding results was improved using BiSCoT v2.3.3 (Istace et al., 2020). A minimum length cutoff of 50 kb was applied to filter out sequences that were not assembled into chromosomes. The naming and orientation of the chromosomes of *R. rugosa* genomes were adjusted to ensure consistency with the published *R. chinensis* (NG) genome. To further scaffold the *R. sericea* genome into chromosome-level pseudomolecules, reference-based scaffolding was conducted using RagTag v2.1.0 (Alonge et al., 2022), with the *R. chinensis* (NG) genome as reference and gap size set to 100 bp.

The completeness of the genomes was evaluated through several approaches. First, HiFi and IsoSeq reads were mapped against the genomes using minimap2 v2.26-r1175 to check the mapping rates. Merqury v1.3 (Rhie et al., 2020) was used to evaluate the base accuracy of the assembled genomes. BUSCO (Manni et al., 2021) was used to further evaluate the completeness (using eudicots odb10), and the tool FCS v0.5.0 was used to check for contamination.

### Genome annotation

Repeat elements were annotated using the EDTA v2.2.0 (Ou et al., 2019) pipeline with parameters set as ‘–anno 1 –sensitive 1’. RepeatModeler v2.0.5 (Flynn et al., 2020) was used to generate the *de novo* repeat libraries for the input genomes. The following tools were used by EDTA to obtain the raw TE candidates: GenomeTools v1.5.10 (Gremme et al., 2013), LTR FINDER parallel v1.1 (Ou and Jiang, 2019), LTR retriever v2.9.5 (Ou and Jiang, 2018), Generic Repeat Finder v1.0 (Shi and Liang, 2019), TIR-Learner v1.19 (Su et al., 2019), HelitronScanner v1.1 (Xiong et al., 2014), and TEsorter v1.4.6 (Zhang et al., 2022). The LTR assembly index (LAI) was calculated using LTR retriever. The repeat annotation results were used to create soft-masked genomes for gene prediction. To reduce over-masking of genic regions, we choose to only mask TEs / repeat elements with lengths over 1 kb.

Genome annotation was performed using a combination of *ab initio* prediction tools and homology-based tools. Helixer v0.3.3 (Holst et al., 2023) was used for *ab initio* gene structural prediction on the non-masked genomes. The following parameters were used: ‘–subsequence-length 106920 –overlap-offset 53460 –overlap-core-length 80190 –lineage land plant’. Additional RNAseq data for *R. rugosa* were obtained from NCBI to assist with annotation (Table S14). First, raw RNAseq reads were filtered using fastp v0.23.3 (Chen et al., 2018) using default parameters. Clean reads were subsequently aligned to the genome assemblies using hisat2 v2.2.1 (Kim et al., 2019) with parameters ‘—dta –max-intronlen 20000’. For the phased *R. rugosa* genome assembly, RNAseq alignments were split into haplotype-specific files based on genomic location, using the samtools split. Braker3 v3.0.8 (Gabriel et al., 2024) was used for gene annotation on the repeat-masked genome. BUSCO with lineage dataset ‘–busco lineage eudicots odb10’ was adopted by Braker3 to evaluate the quality of the intermediate annotations. For homology-based predictions, the Viridiplantae sequences from OrthoDB v11 (Kuznetsov et al., 2023), together with the protein sequences of *R. chinensis* and *R. rugosa* from the NCBI RefSeq database were used. Miniprot v0.11 (Li, 2023) was used to generate homology predictions. The Isoseq data were processed using IsoSeq v3.8.2. After quality control and clustering, the reads were aligned to the assemblies using minimap2 ‘-ax splice:hq –uf’. Redundant alignments were removed, and the Braker3 long-read pipeline was employed to predict gene structures from the IsoSeq alignments. EVidenceModeler v2.1.0 (Haas et al., 2008) was used to integrate the *de novo* prediction, protein alignment and transcript prediction results. The following weights were used: 1 for *ab initio* predictions, 5 for protein alignment predictions, and 10 for transcript predictions. PASA v2.5.3 (Haas et al., 2008) was used to refine the EVidenceModeler predictions with the information on the untranslated regions (UTRs) and alternative splicing (AS).

The PASA annotations were further validated using AGAT v1.4.0 (Dainat et al., 2024) and Gff3toolkit v2.1.0 (https://github.com/NAL-i5K/GFF3toolkit). Annotations with errors including coding sequence phase issues, incomplete gene structures, gene overlaps, or abnormal gene lengths (e.g. *<*100 kb) were either manually corrected or removed. Single-exon genes were filtered using a hierarchical decision tree incorporating four evidence types: evolutionary conservation (BUSCO against eudicots odb10), RNA-seq expression support (≥10X read depth and ≥70% transcript coverage across mapped reads), functional domain annotation (InterProScan v5.67-99.0 (Jones et al., 2014)), and homology grouping derived from PanTools v4.3.1 (Jonkheer et al., 2022) using available *Rosa* genome annotations (Fig. S27). Homology groups were classified into three tiers: (1) genes grouped with RefSeq annotations, (2) genes grouped with all non-RefSeq *Rosa* annotations, and (3) genes grouped with part of non-RefSeq *Rosa* annotations. The decision tree prioritized genes meeting BUSCO completeness thresholds, followed by those with RNA-seq expression evidence, InterPro domain matches, and homology relations. Genes failing all criteria were excluded to minimize false-positive annotations. Functional annotations were added to the gff3 files through the steps below: First, the reciprocal best hits based on two RefSeq rose annotations (GCF_002994745.2 and GCF_958449725.1) were added; Next, PFAM domain annotation results obtained from InterProScan were also added to the gff3 files using AGAT; Remaining transcripts lacking annotation were labeled “hypothetical protein”. The annotation of the *R. sericea* genome was transferred to the reference-based chromosome-level assembly by RagTag.

### Rosaceae panproteome

The Rosaceae proteomes were clustered into homology groups (HGs) using PanTools in relaxation mode six based on the optimal grouping procedure. Intersections of the homology groups across the Rosaceae panproteome were visualized using ComplexUpset v1.3.5 (Krassowski, 2020). To build the species phylogenetic tree, we first generated the protein alignment for each core HG using MAFFT v7.520 with parameter ‘–auto’ (Katoh et al., 2002). Subsequently, protein alignments were converted to nucleotide alignments with PAL2NAL v14 (Suyama et al., 2006) and trimmed with trimal v1.4.1 (Capella-Gutiérrez et al., 2009). Finally, IQ-tree v1.6.12 (Minh et al., 2020) and ASTRAL-Pro v1.16.2.4 (Zhang et al., 2020) were applied for species tree construction. The mcmctree from PAML v4.10.7 (Yang, 2007) was used to estimate the species divergence times. The JC69 model and independent evolution rates were selected for mcmctree. The following time calibrations were used: 90 Mya as the minimum age for the Rosaceae crown; 55 Mya as the minimum age for divergence between Maleae and Amygdaleae; 47.8 Mya as the minimum age for divergence between Roseae and other clades in Rosaceae (Li et al., 2011; Xiang et al., 2016; Zhang et al., 2017). Contraction and expansion of the HGs in Rosaceae was modeled using CAFE5 v1.1 (Mendes et al., 2021) using the gamma model and parameter ‘k=3’ to account for rate variation among homology groups. HGs containing members from only one species or those with more than 100 members from a single species were excluded from the CAFE analyses. Functional annotations of the Rosaceae panproteomes were obtained by searching the eggNOG v5.0 database using eggNOG-mapper v2 (Huerta-Cepas et al., 2019; Cantalapiedra et al., 2021). TBtools-II v2.148 (Chen et al., 2023) was used for KEGG and GO enrichment analysis. We filtered the GO terms based on Benjamini-Hochberg (FDR)-adjusted p-value using 0.05 as threshold. For KEGG and GO enrichment of branch-specific homology groups, we used branch-specific lists of homology groups under expansion or contraction as query sets, with branch-specific backgrounds comprising existing homology groups at each branch that had GO annotations.

### *Rosa* Pangenome analyses

Publicly available *Rosa* assemblies and annotations were collected for building the pangenome (Table S3). Genomic sequences shorter than 5 kb and genes with less than 30 amino acids were excluded by PanUtils QC pipeline (https://github.com/PanUtils/pantools-qc-pipeline) to ensure high data quality (Table S15). Mash v2.3 was used to estimate the genetic distance among the *Rosa* genomes (Ondov et al., 2016). PanTools v4.3.1 was used to construct the *Rosa* pangenome. Macrosynteny across the haploid *Rosa* genomes was analyzed using GENESPACE v1.2.3 (Lovell et al., 2022). We adapted our criteria for defining core, accessory, and unique HGs to the species level to account for cases where multiple genomes from the same species were included in the pangenome. To identify core HGs with species- or clade-specific expansion or contraction, we first normalized HG counts across phased genomes to ensure a consistent haplotype-level comparison. Then, we calculated the fold change between two groups for each HG. HGs with fold change greater than 1.5 were kept for further KEGG and GO functional enrichment analyses with same procedure applied in Rosaceae panproteome analyses.

### PanTE analysis

The panEDTA module from EDTA (Ou et al., 2024) was used for analyzing the TEs across the *Rosa* pangenome. The coding sequences of *R. chinensis* (NG) were used as input reference for the panEDTA analysis. Full-length LTR insertion time was estimated using T = *K*/2, where *K* is the divergence rate and is the neutral mutation rate (Bowen and McDonald, 2001). *K* was estimated using the Jukes-Cantor model for non-coding sequences with *K* = −3*/*4*ln*(1 − 4*/*3*d*) (Jukes and Cantor, 1969), where d = 100% - identity% is the proportion of sequence differences (Ou and Jiang, 2018).

### Region of interest (ROI) analysis of *Rosa* pangenome

To build the phylogeny for the *Rosa NUDX1* genes, we applied the same methodology used for species tree construction in the Rosaceae panproteome analysis. The *NUDX1* gene (GenBank: NP_177044) from the *A. thaliana* proteome was used as an outgroup (Yao et al., 2017). Ggtree v3.10.1 was used for phylogenetic tree visualization (Yu et al., 2017). The Conserved Domain Database (CDD) was used to identify the protein domains in *NUDX1* genes (Yang et al., 2020). Gene structure and protein domains of NUDX1 were visualized using TBtools-II v2.148. We used PanTools’ blast and extract_region (commit 107838f) functions to identify and extract the region of interest from the pangenome using *R. chinensis* (NG) *NUDX1* family as query. The extracted regions containing members from *NUDX1* family were visualized using gggenomes v1.0.0 (Hackl et al., 2024). MAFFT v7.520 was used to align the identified LINE type TEs. Jalview v2.11.4.0 was used to visualize the multiple sequence alignments (Waterhouse et al., 2009). MEME Suite v5.5.7 was used to identify motifs (Bailey et al., 2015). The identified motifs were compared against the JASPAR2022 CORE database (Castro-Mondragon et al., 2022) by the Tomtom tool from the MEME Suite.

## Availability of data and materials

Genome sequences and gene annotation data for this project were deposited at NCBI under BioProject accession numbers: PRJNA1091549 (hapA of *R. rugosa*), PRJNA1091550 (hapB of *R. rugosa*), PRJNA1117493 (*R. sericea*). The Bionano maps and raw sequencing reads, including the genomic HiFi data and RNA-seq data, have been deposited under the corresponding NCBI BioProject accessions listed above. All commands needed to reproduce the pangenome can be found in Supplementary file *reproduce_pangenome*.*md*.

## Acknowledgements

We gratefully acknowledge Mirjam Lemmens, collection manager at the Belmonte Arboretum Foundation, Wageningen, for her invaluable assistance and advice with sampling of the *Rosa* species, and we thank Erik Wijnker, Sara Diaz Trivino, Freek Bakker, Judith Risse, and Sander Peters for their contributions to generating the genomic resources and preliminary data analysis. We also thank Christina Papastolopoulou for her expert advice on genome assembly and annotation, and Dirk-Jan van Workum for his support with PanTools.

