## Supplemental Figures for "Two high-quality rose genomes underpin a novel *Rosa* pangenome to advance rose genomics, phylogenetics, and breeding"

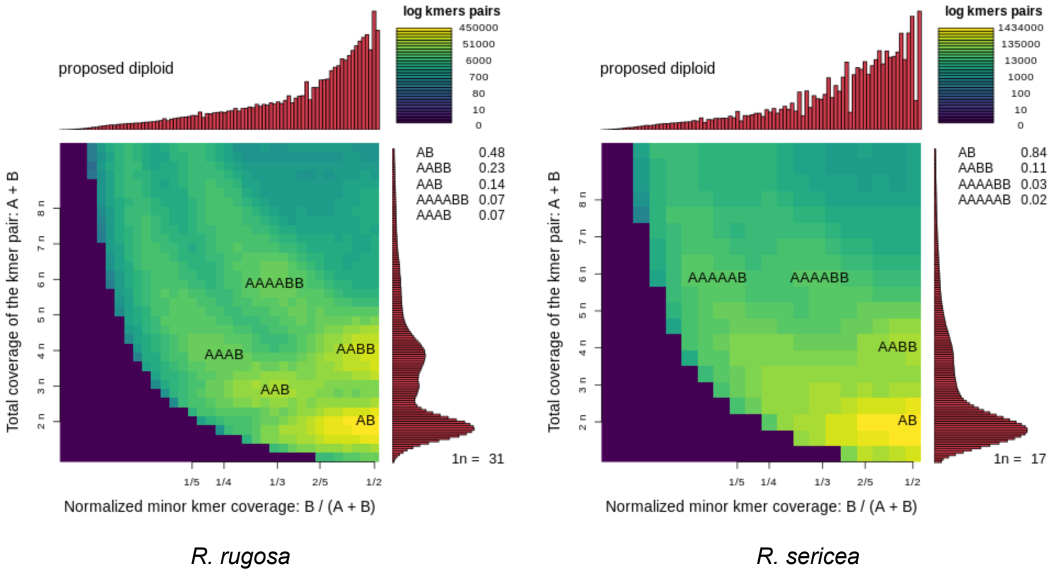


Figure S1: Smudgeplot analyses of two *Rosa* species for ploidy level estimation.

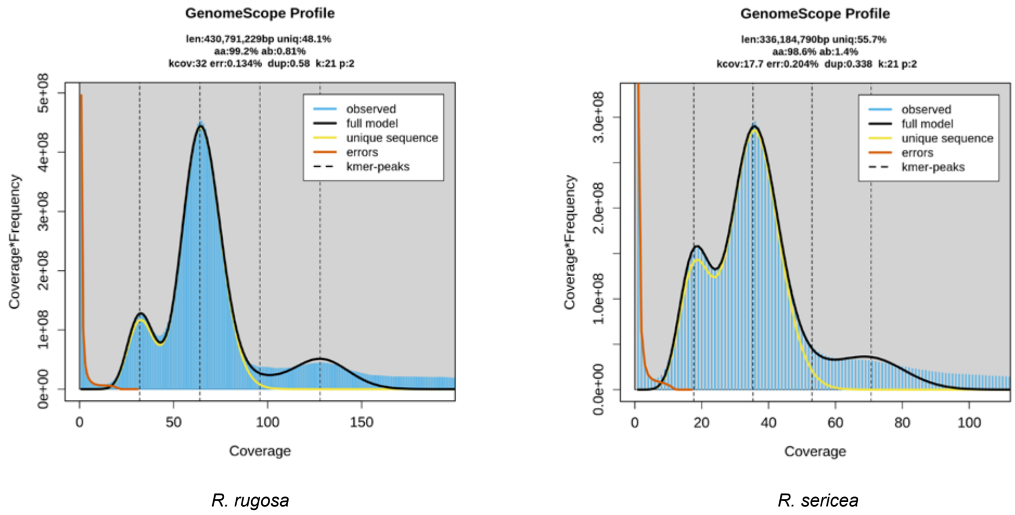


Figure S2: GenomeScope2 analyses of two *Rosa* species using 21-mer frequency spectra.

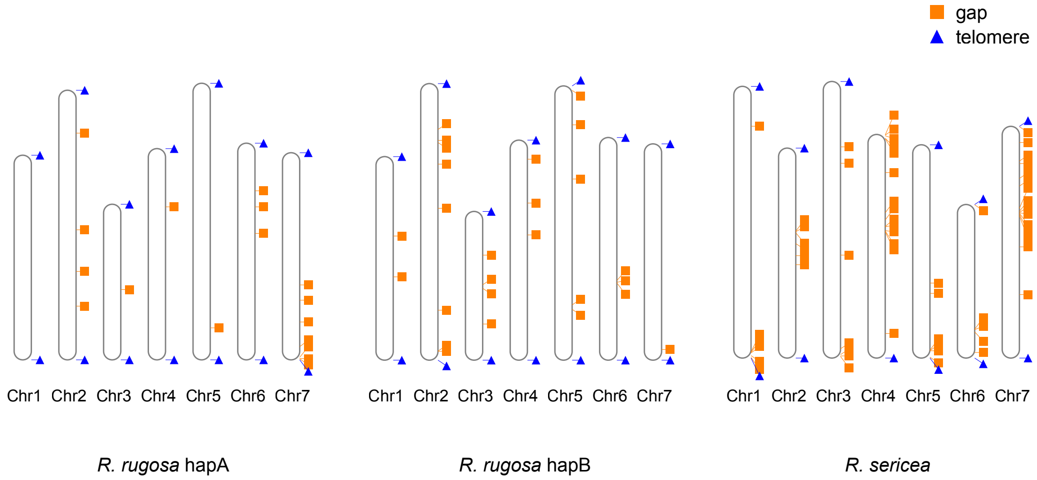


Figure S3: Telomeres and assembly gaps in *R. rugosa* (hapA: 17 gaps; hapB: 26 gaps) and *R. sericea* (59 gaps) genomes. Triangles and boxes only indicate positions (to scale).

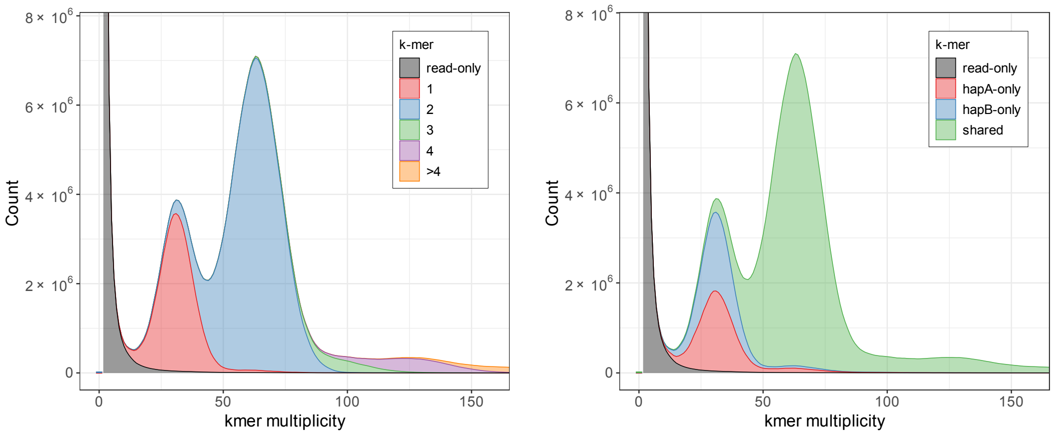


Figure S4: Merqury analyses of the haplotype-resolved *R. rugosa* genome. The left plot shows the copy number spectrum of k-mers colored by the copy numbers found in the assembly. The right plot shows distinct k-mer assembly spectrum with assembly-specific k-mers (red and blue) and shared portion of k-mers (green)

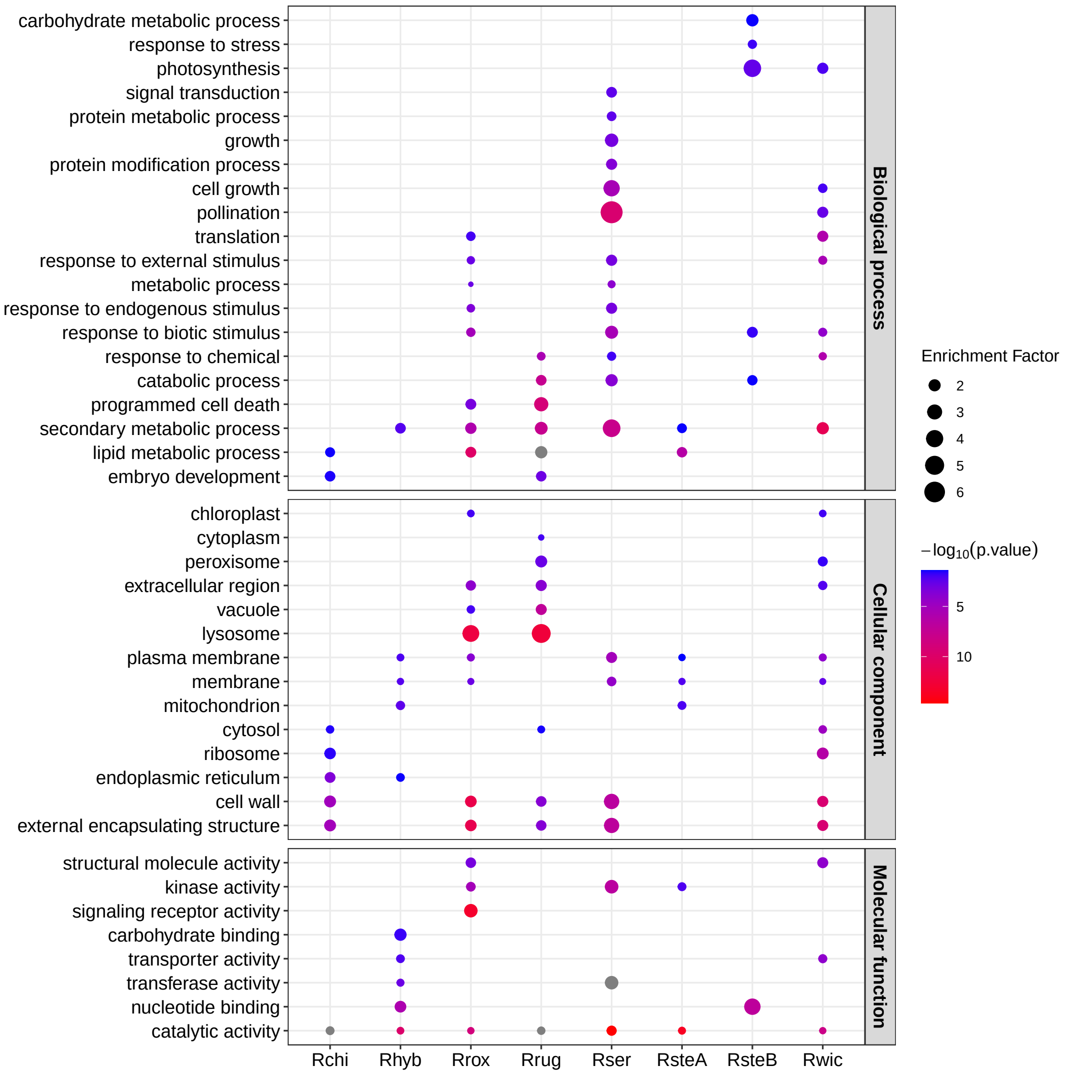


Figure S5: GO enrichment analysis of homology groups under significant expansion in sampled *Rosa* genomes.

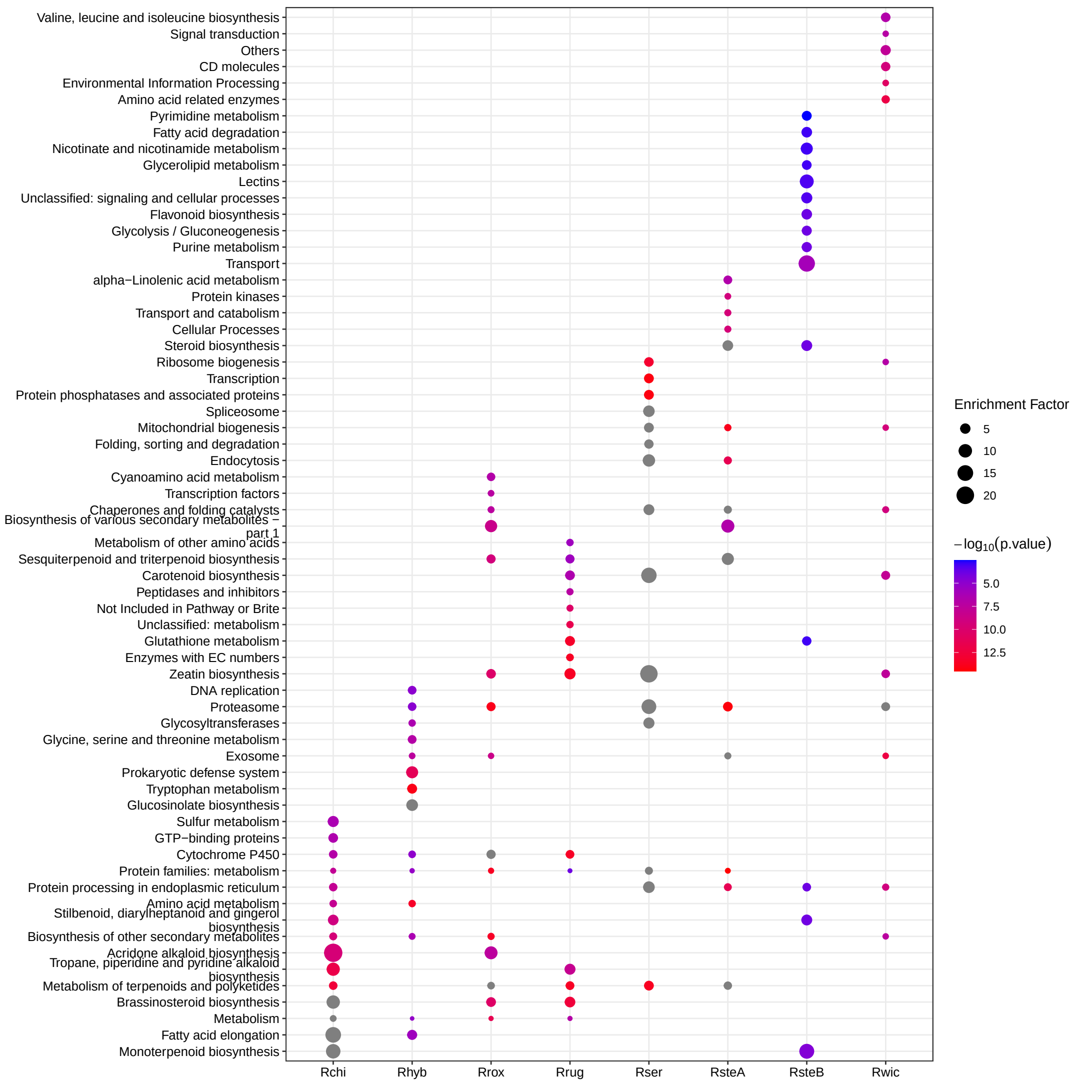


Figure S6: KEGG enrichment analysis of homology groups under significant expansion in sampled *Rosa* genomes.

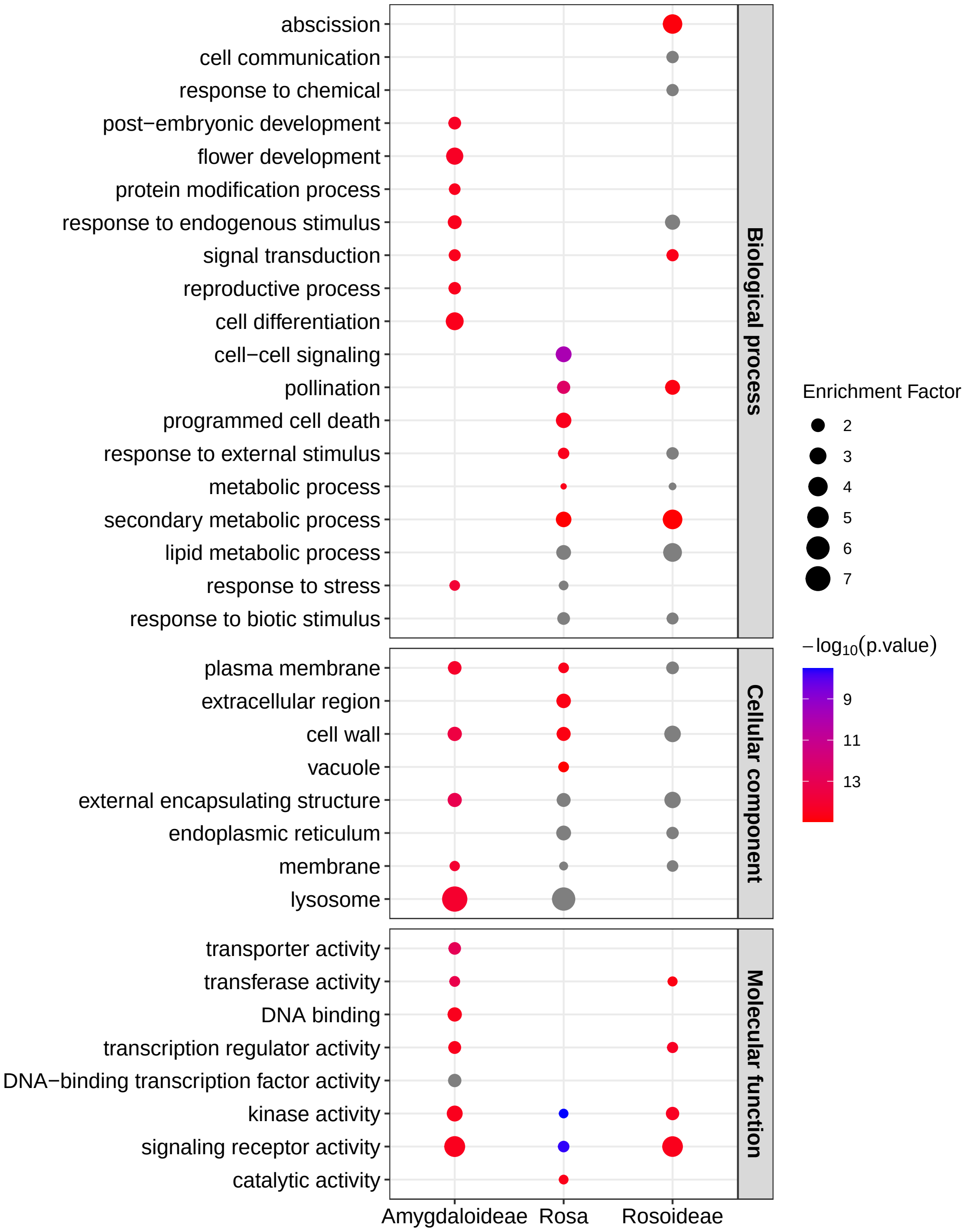


Figure S7: GO enrichment analysis of homology groups under significant expansion in Amygdaloideae, *Rosa*, and Rosoideae.

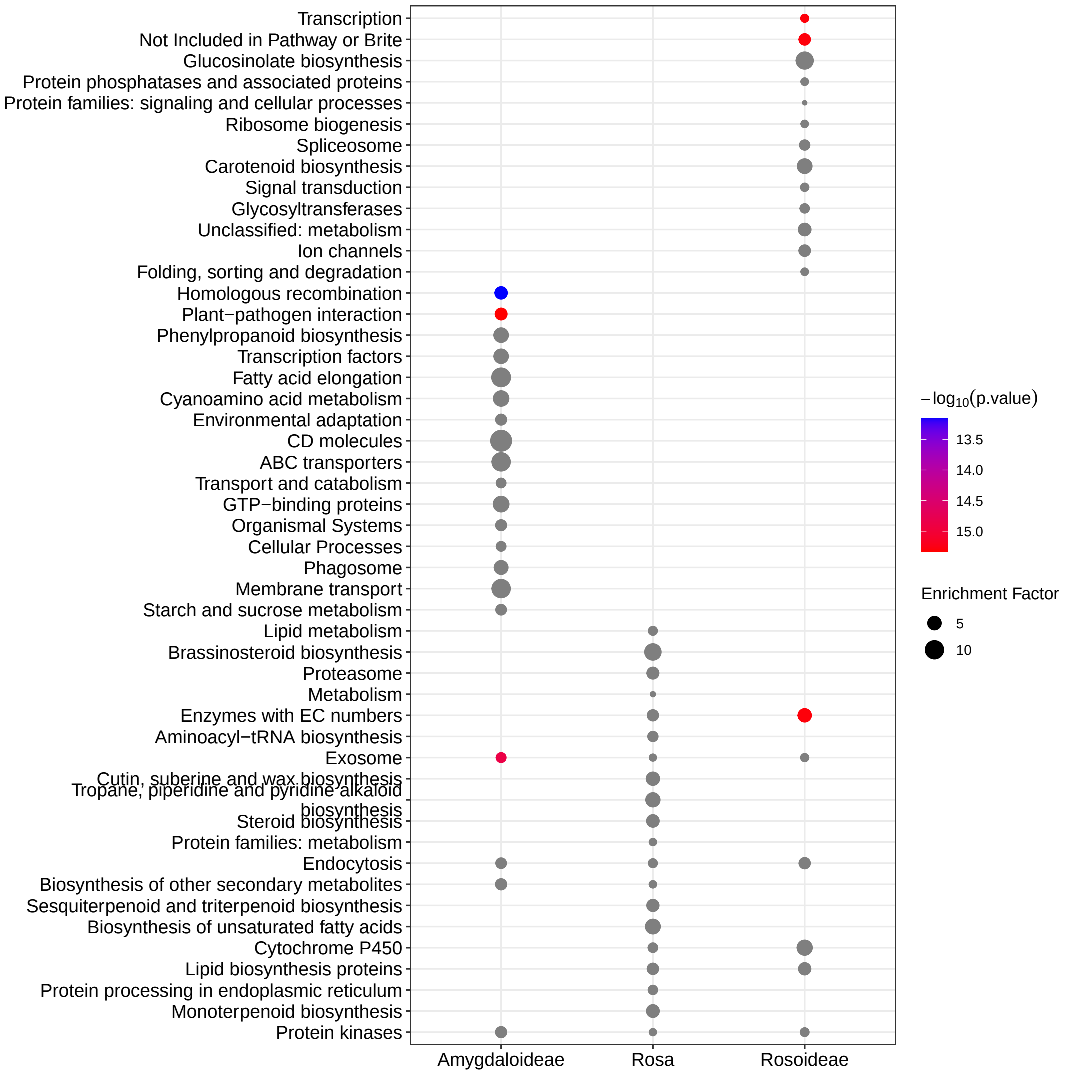


Figure S8: KEGG enrichment analysis of homology groups under significant expansion in Amygdaloideae, *Rosa*, and Rosoideae.

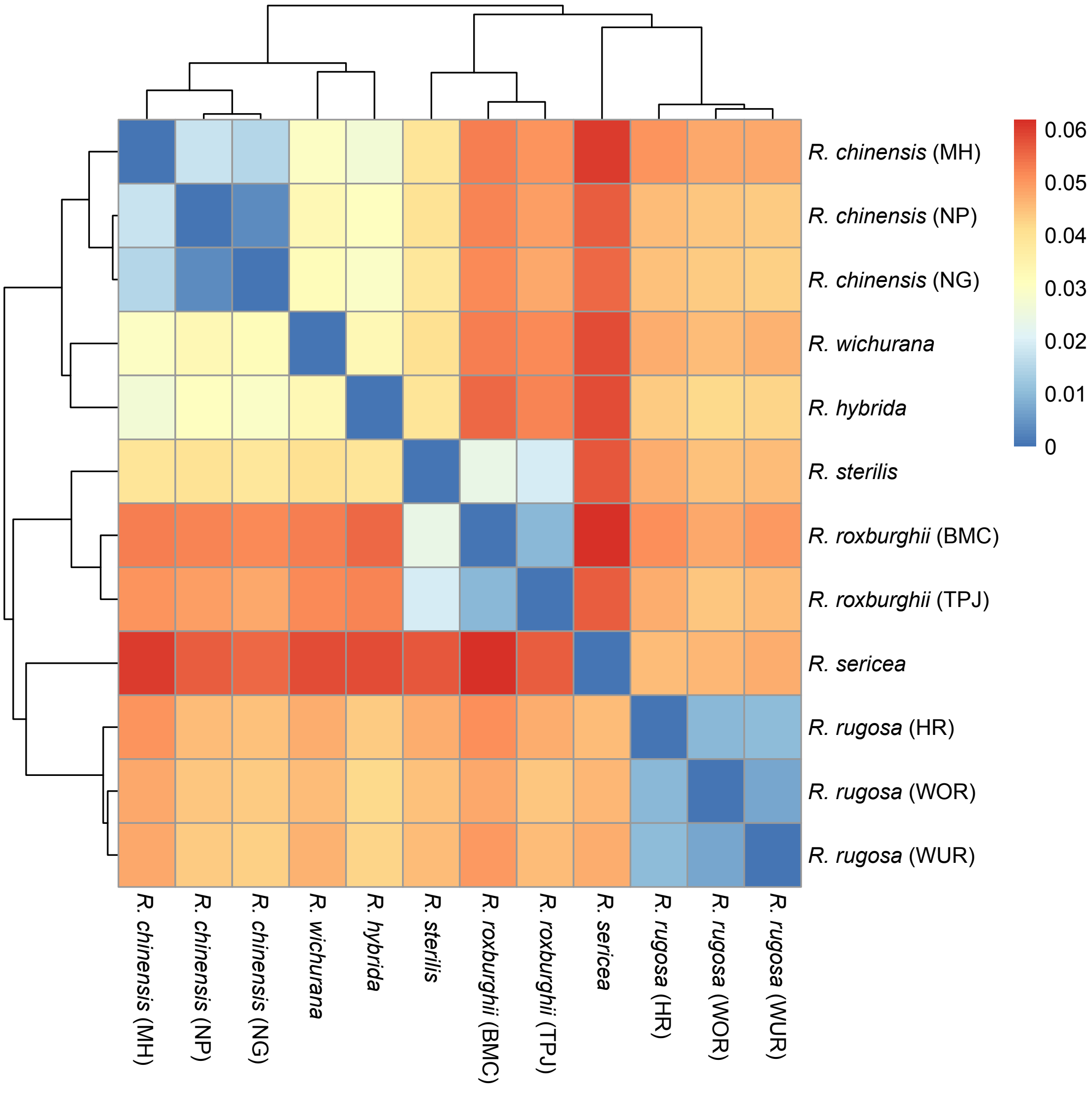


Figure S9: Mash distance of the *Rosa* genomes, used for pangenome construction.

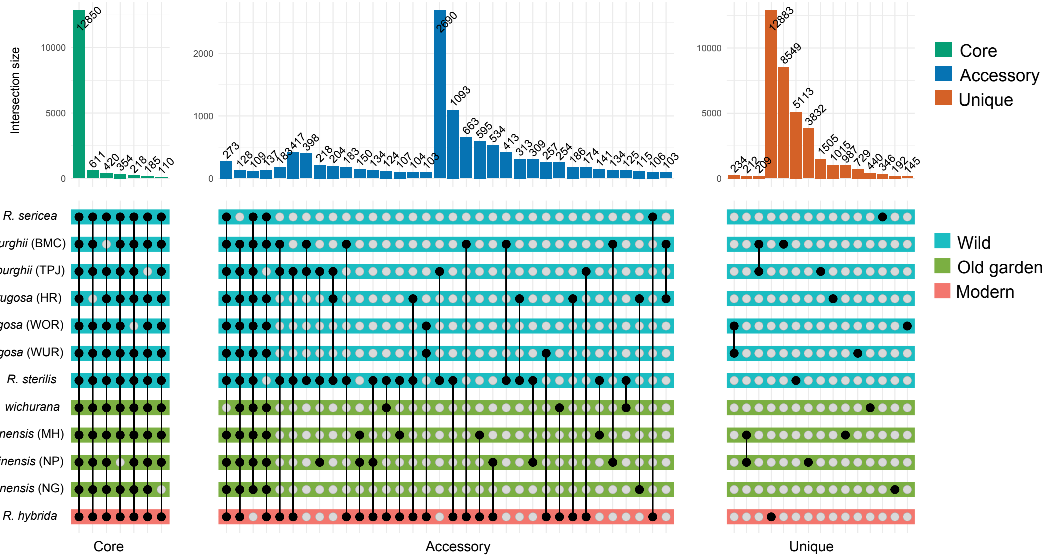


Figure S10: Upset plot of core, accessory, and unique homology groups from *Rosa* pangenomes. The intersections were sorted by degree for the three types of homology groups. Intersections with size larger than 100 were shown.

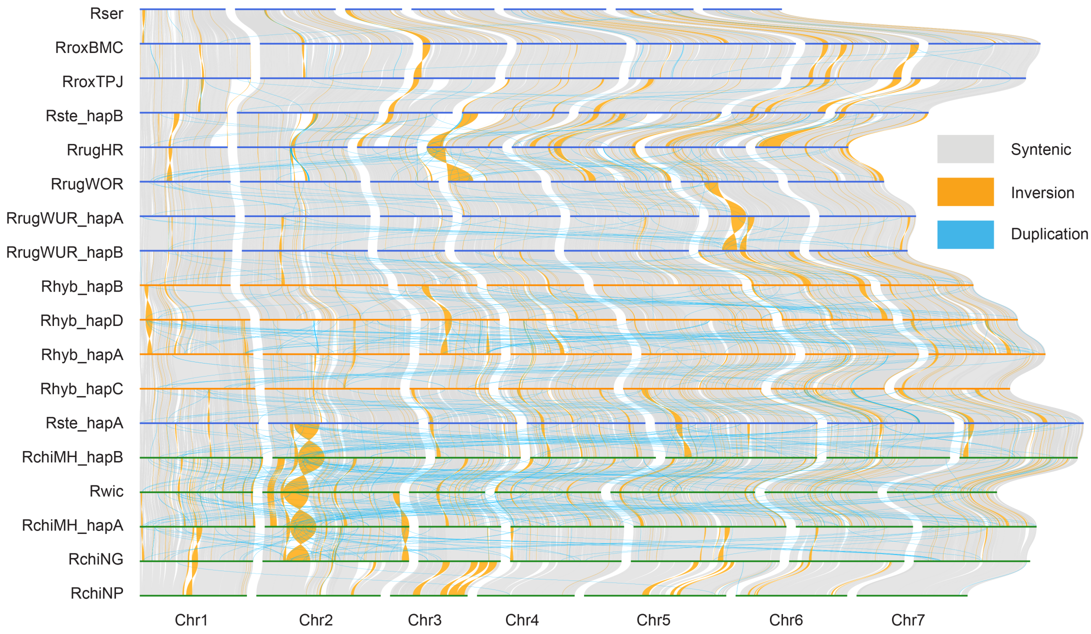


Figure S11: Landscape of sequence divergence across the *Rosa* pangenome.

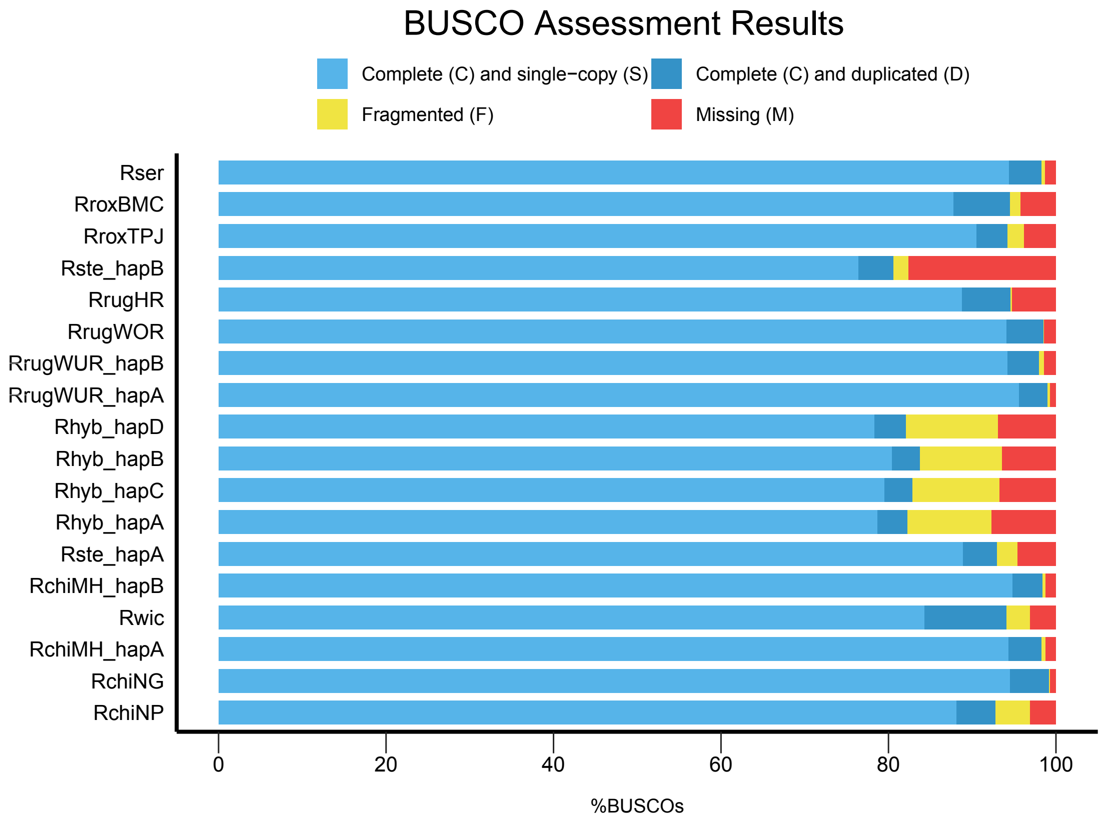


Figure S12: BUSCO evaluation of the input *Rosa* genomes.

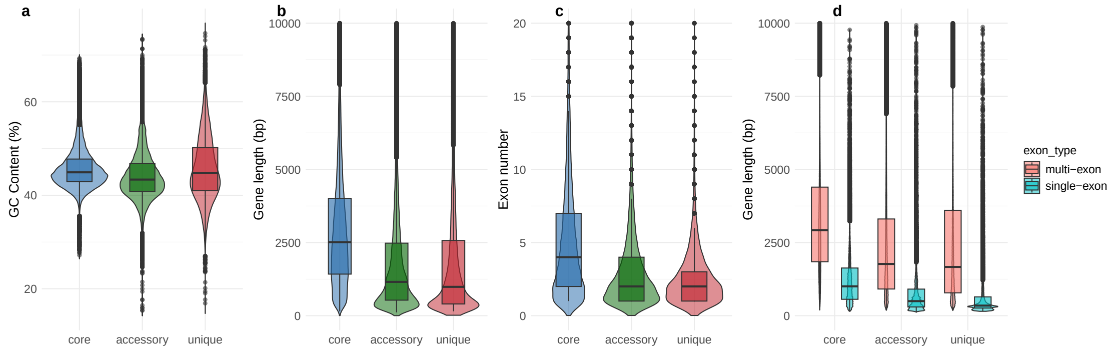


Figure S13: Metrics for core, accessory, and unique genes in *Rosa* pangenome. **a** GC content. **b** Gene length. **c** Exon number. **d** Gene length of single-exon and multi-exon genes.

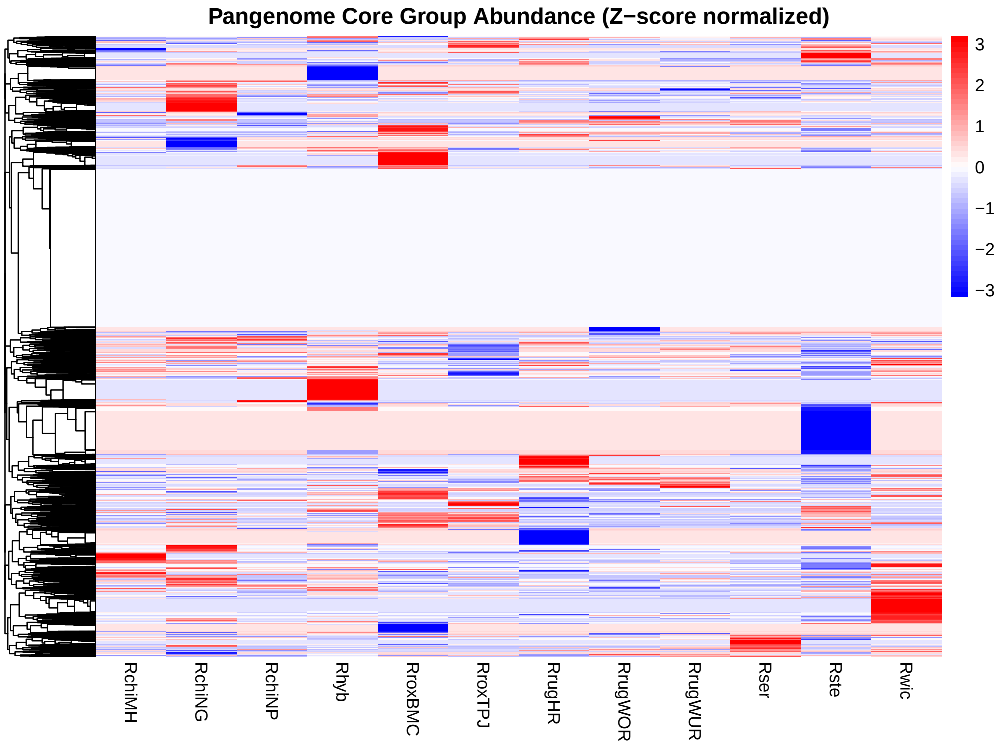


Figure S14: Abundance of core homology groups of *Rosa* pangenomes. Gene counts for each homology group were normalized using Z-score. Blank indicate homology groups without variations.

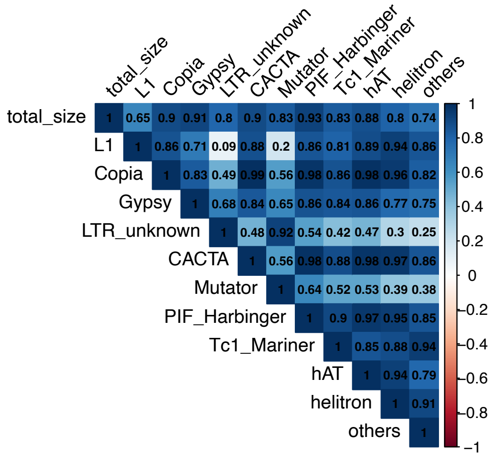


Figure S15: Correlation of different types of TE and genome size in the *Rosa* pangenome

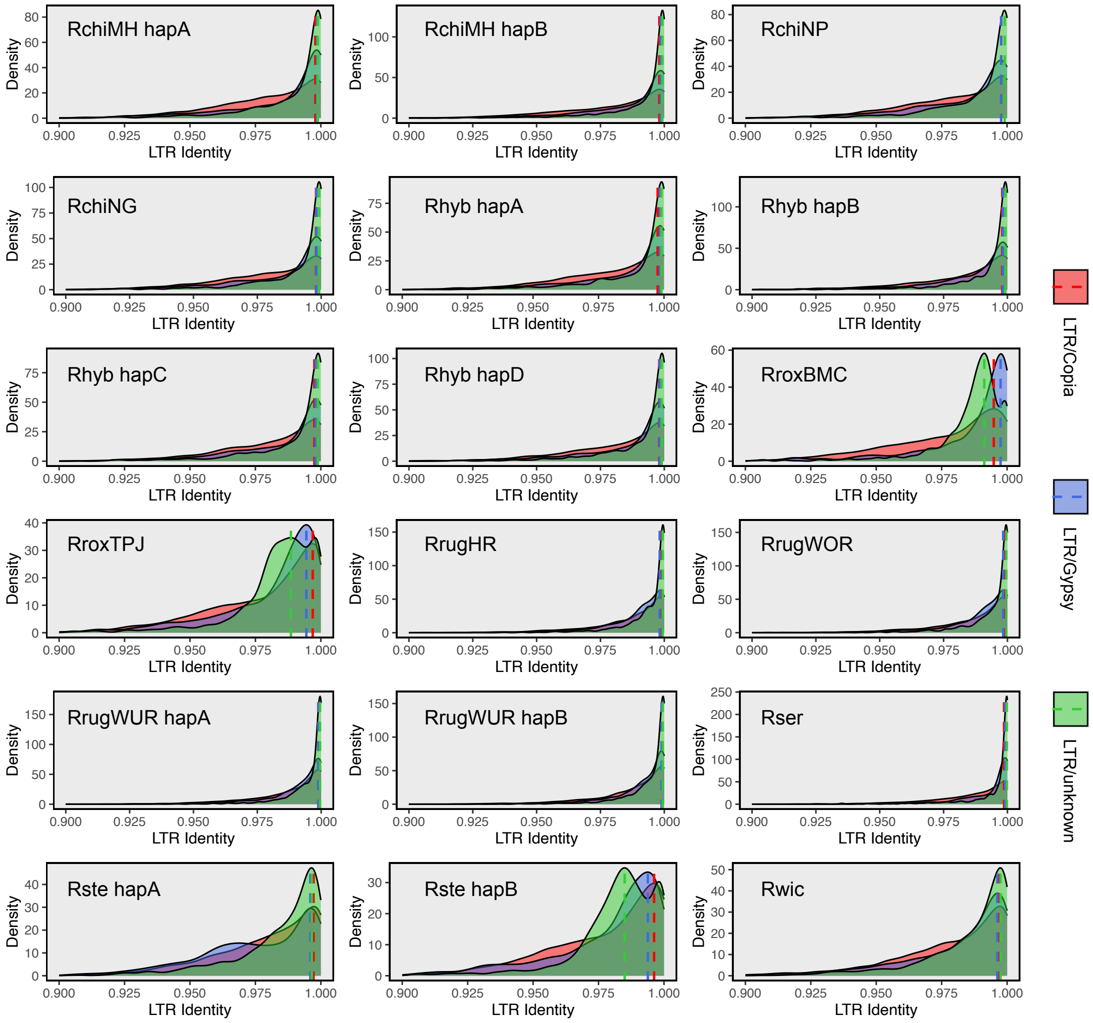


Figure S16: Distribution of intact LTRs identity for different LTR classes in *Rosa* genomes. The vertical dashed lines indicate peaks of density distribution.

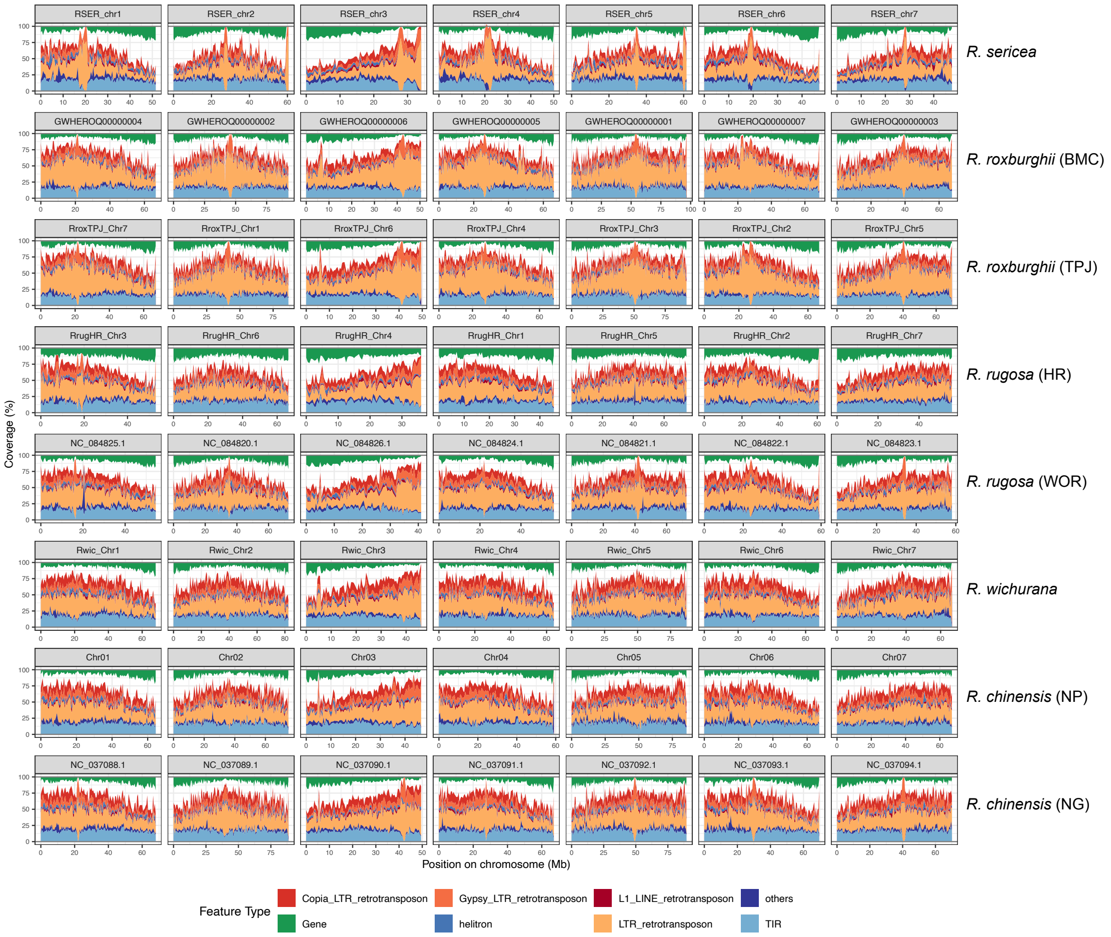


Figure S17: Chromosomal gene and repeat density for non-haplotype-resolved *Rosa* genomes.

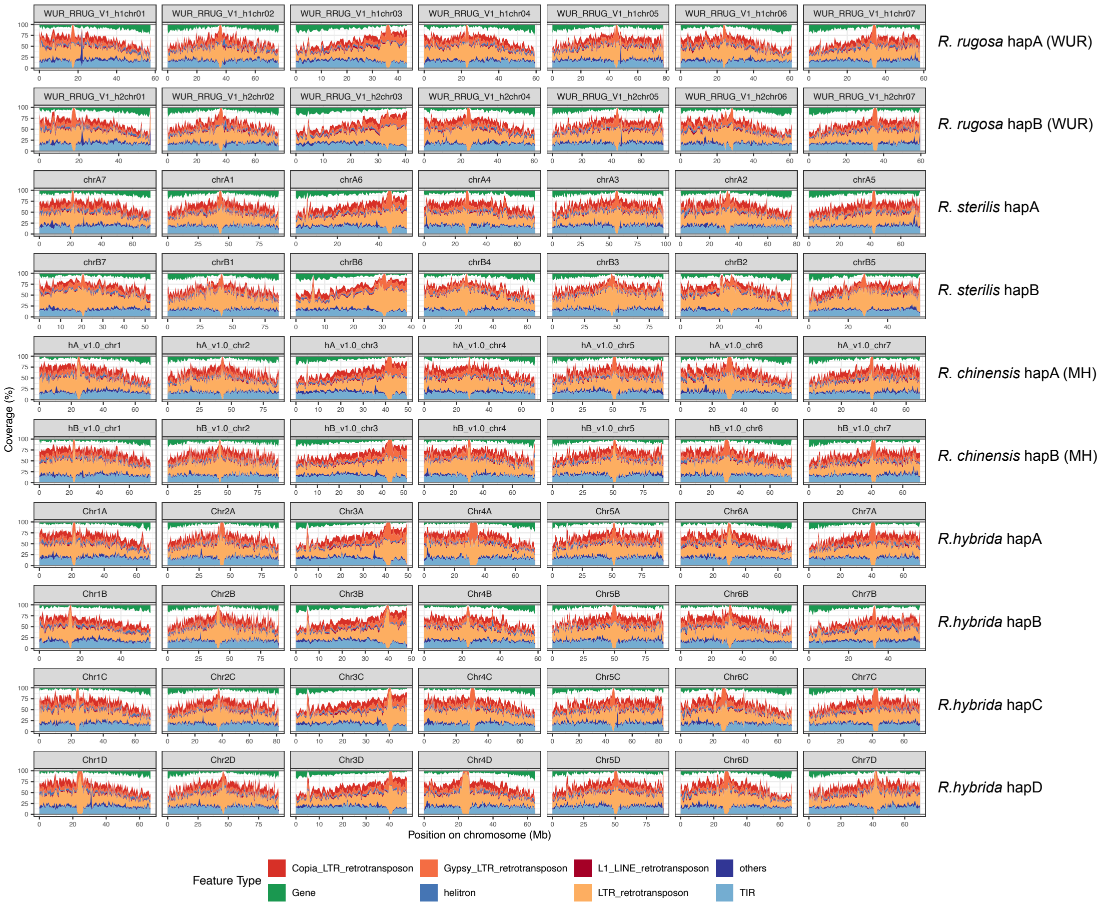


Figure S18: Chromosomal gene and repeat density for haplotype-resolved *Rosa* genomes.

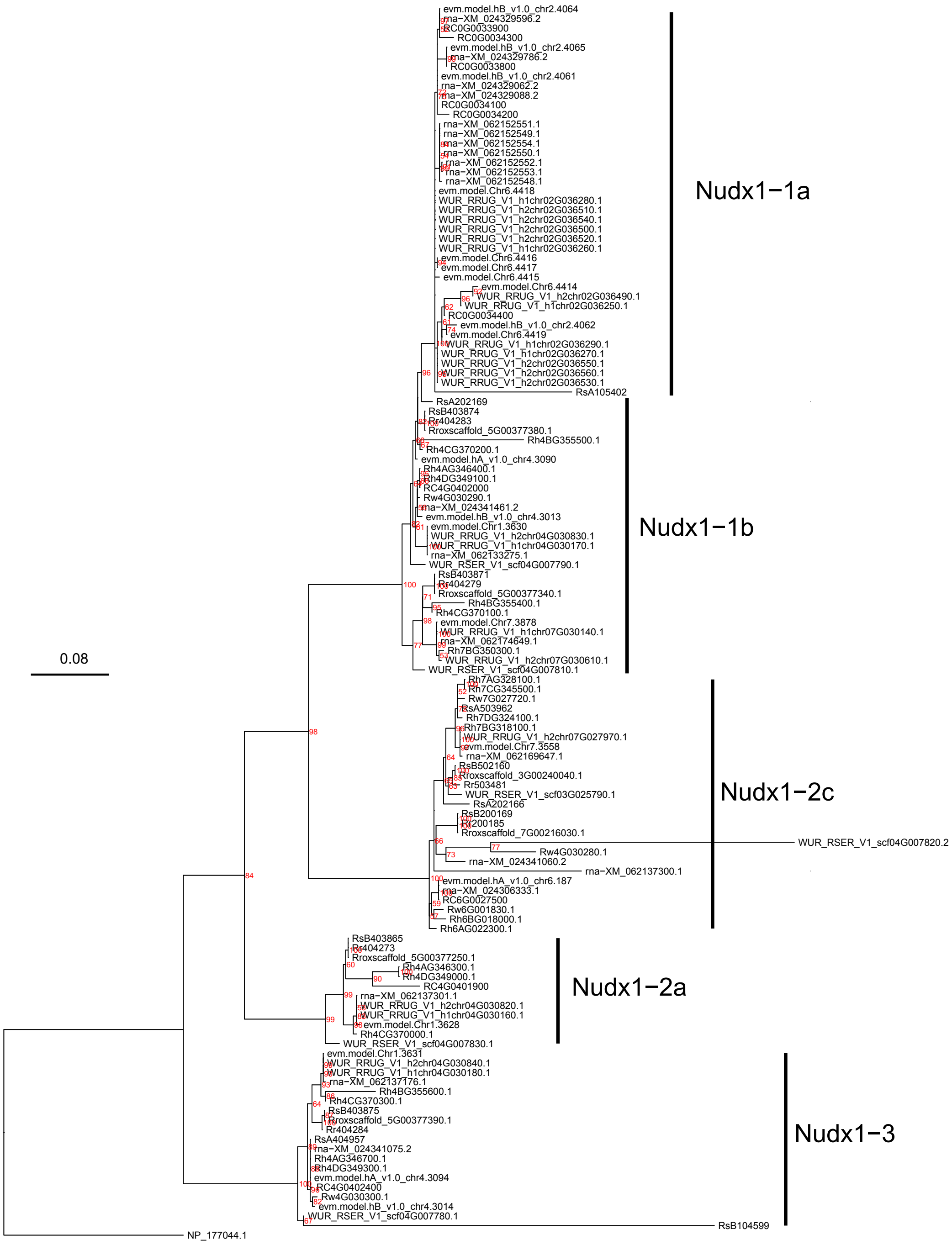


Figure S19: Phylogenetic analysis of the *Rosa* NUDX1 gene family. The red numbers indicate bootstrap support of 1,000 replicates.

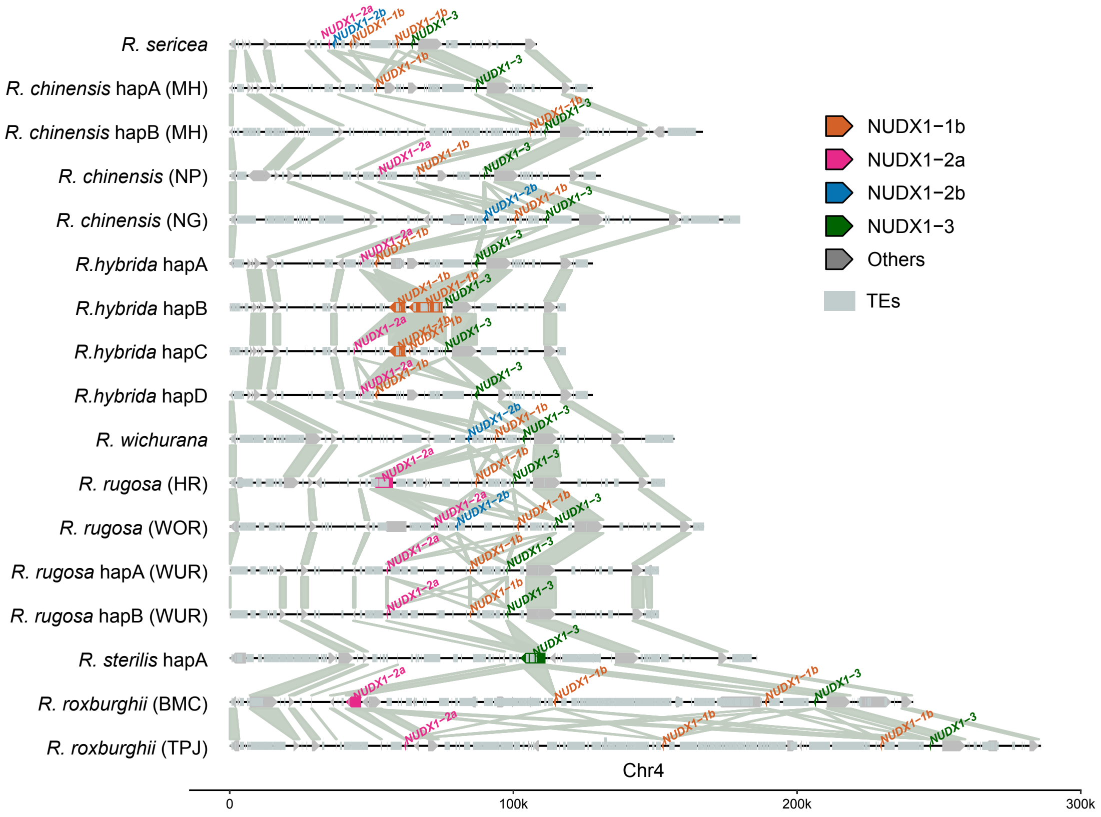


Figure S20: Genomic organization of the NUDX1 gene family region on Chr4 across the *Rosa* pangenome.

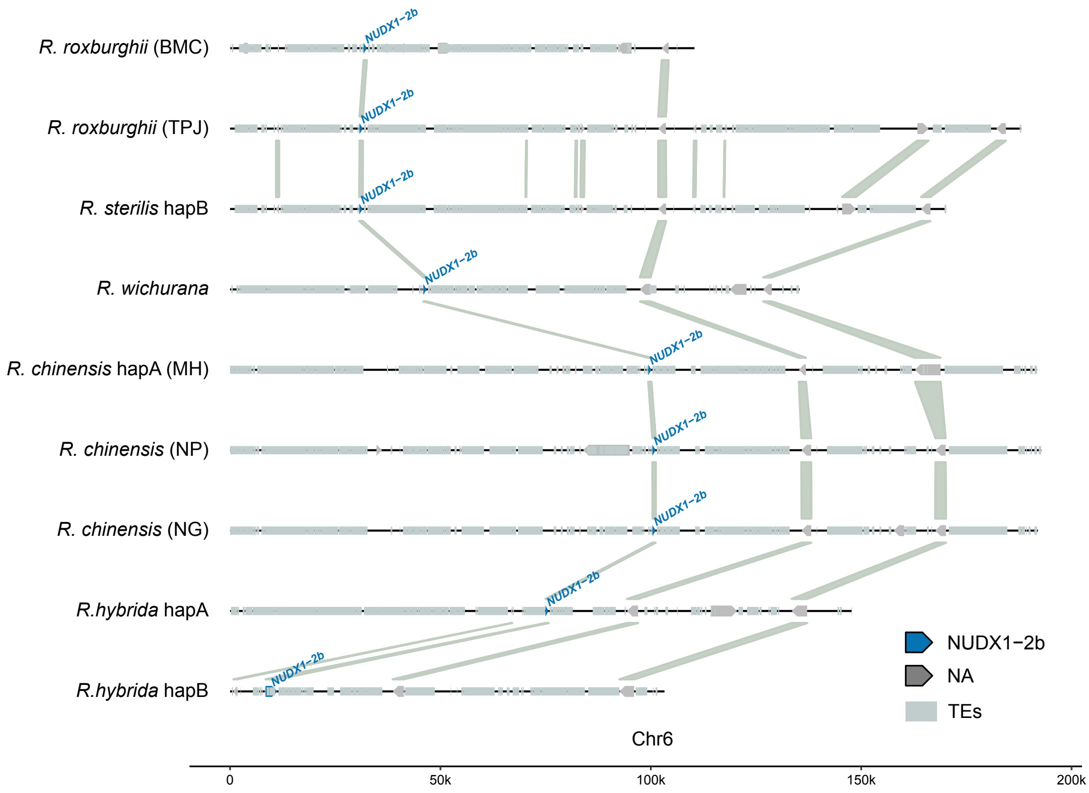


Figure S21: Genomic organization of the NUDX1 gene family region on Chr6 across the *Rosa* pangenome.

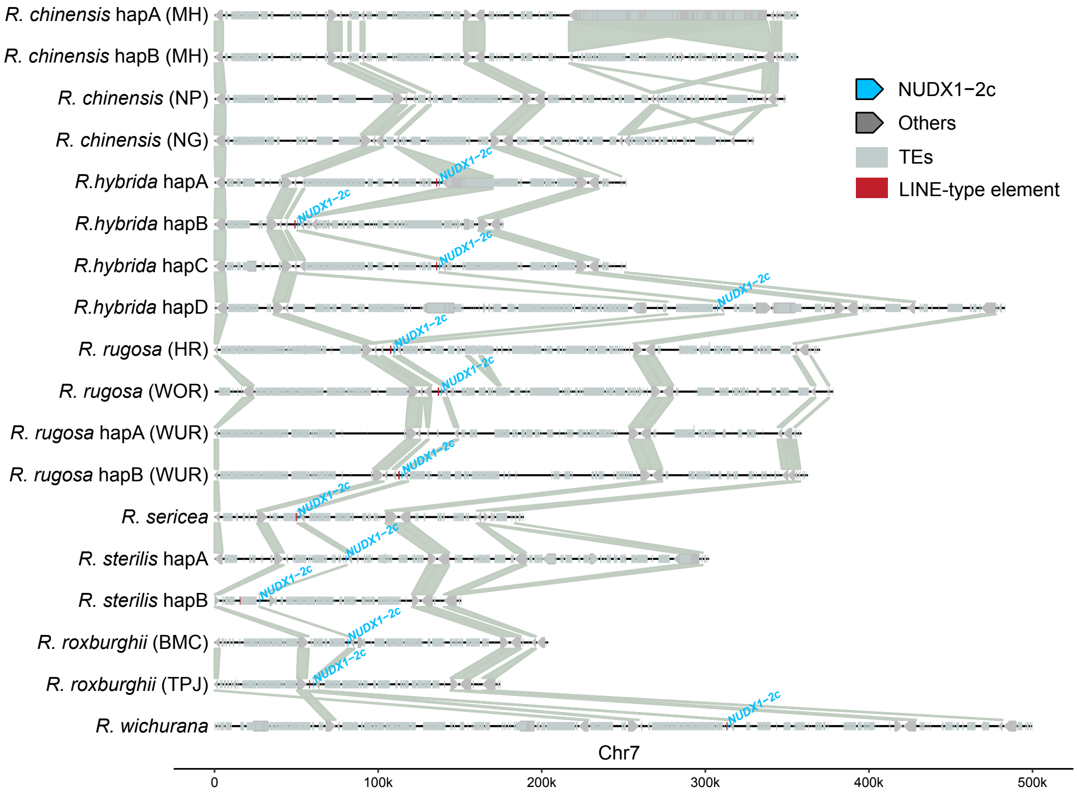


Figure S22: Genomic organization of the NUDX1 gene family region on Chr7 across the *Rosa* pangenome.

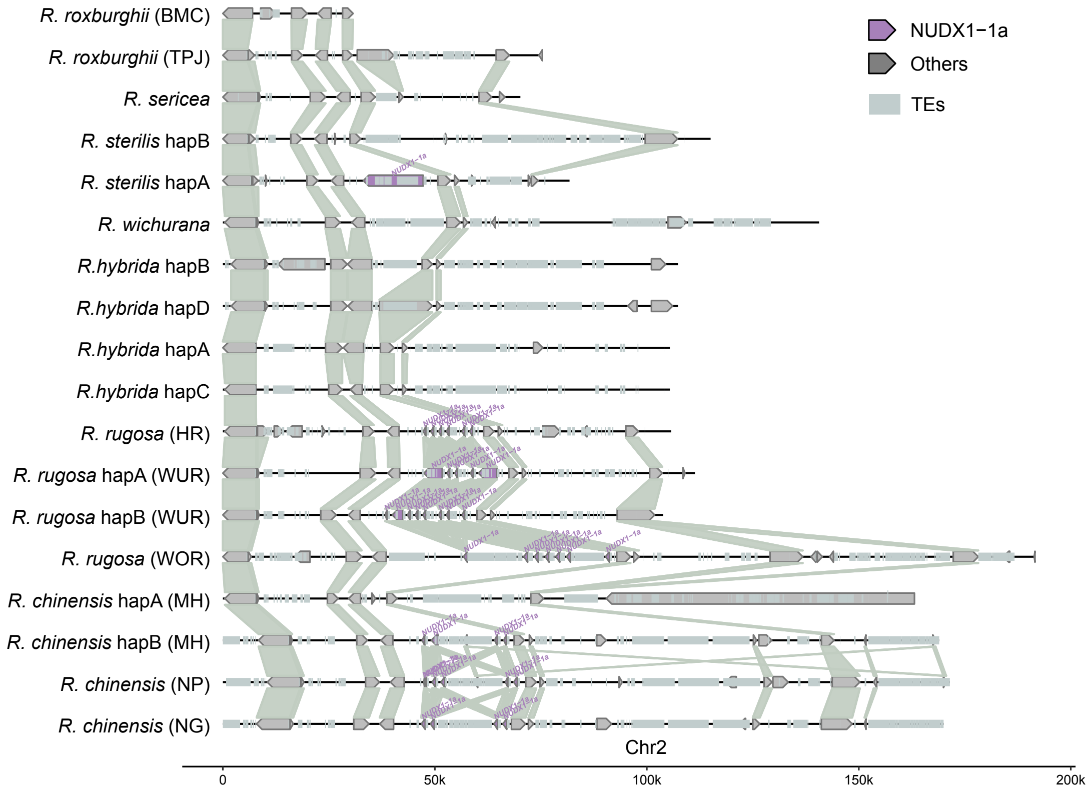


Figure S23: Genomic organization of the NUDX1 gene family region on Chr2 across the *Rosa* pangenome.

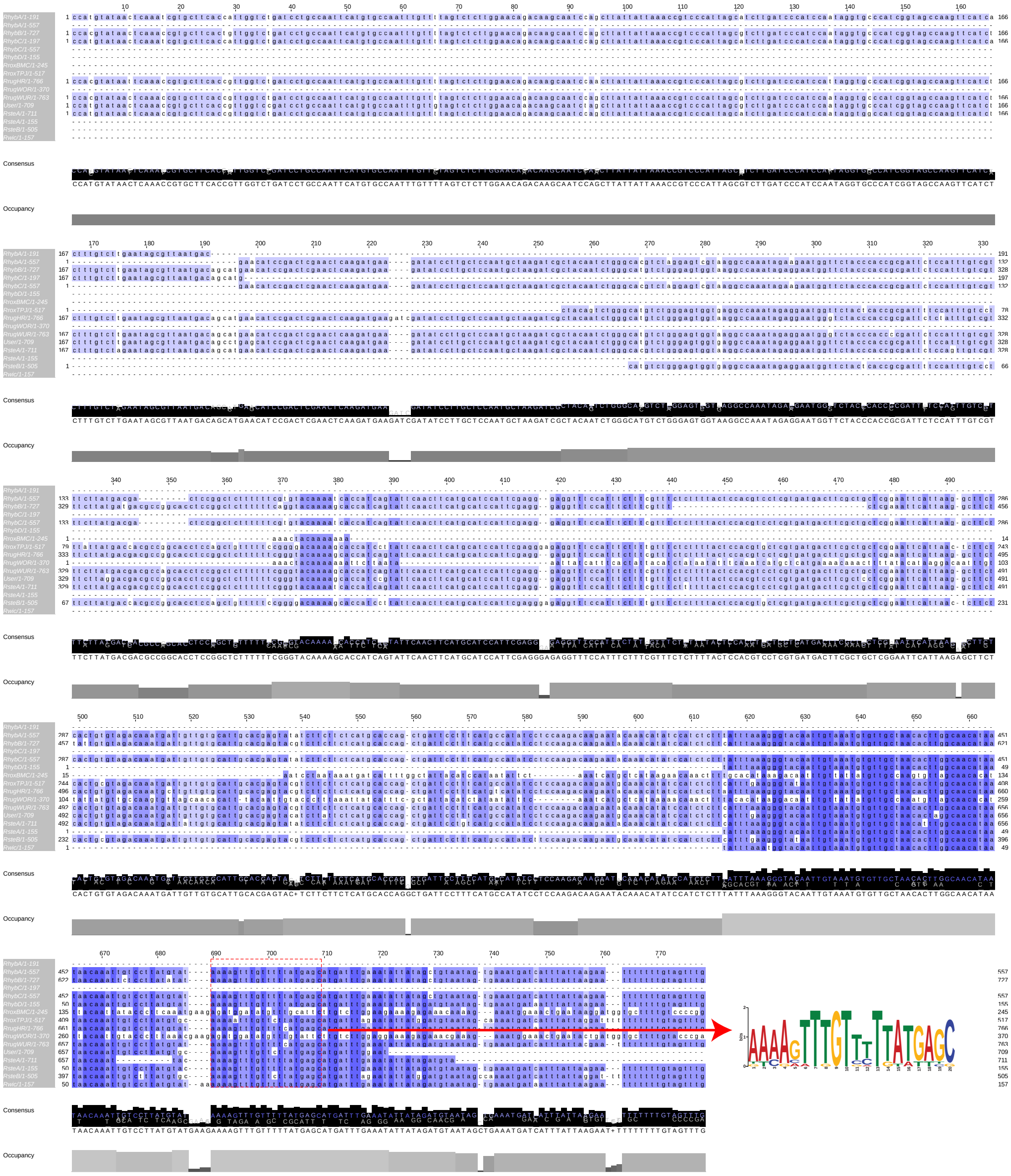

Figure S24: Multiple sequence alignment of LINE type TEs in the upstream of NUDX1-2c genes on Chr7. The highlighted motif represents predicted transcription factor binding site belonging to C2H2 zinc finger factors.

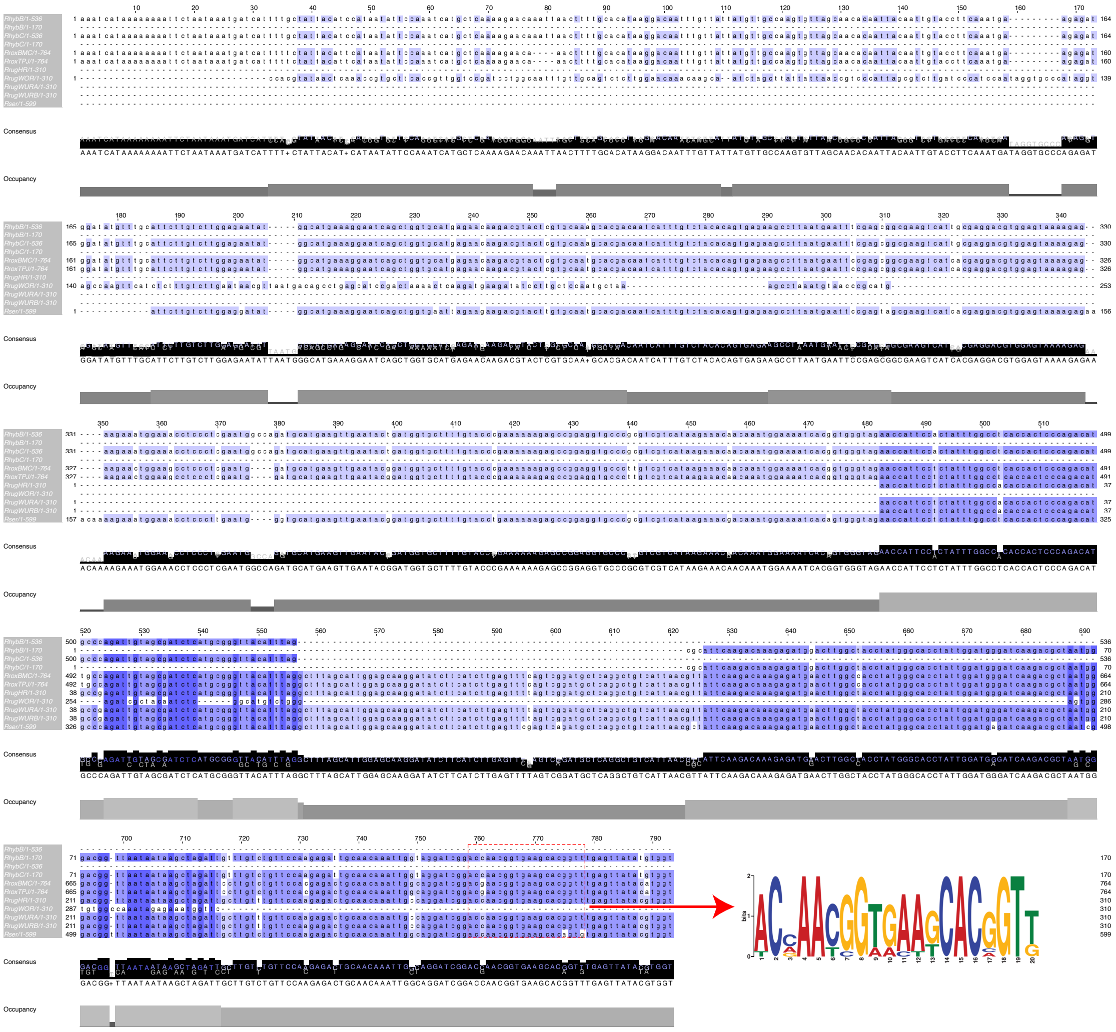

Figure S25: Multiple sequence alignment of LINE type TEs within the tandem array of NUDX1 genes on Chr4. The highlighted motif represents predicted transcription factor binding site belonging to Tryptophan cluster factors.

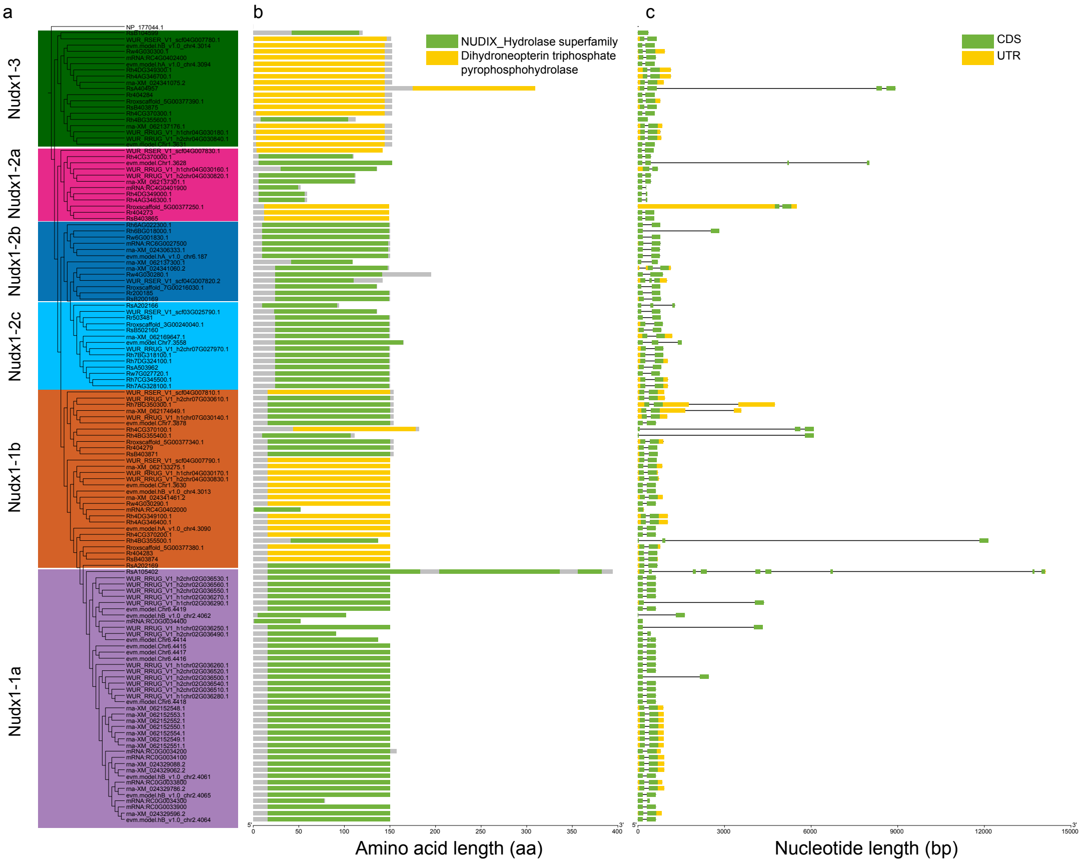


Figure S26: Gene structure and domain analyses of *Rosa* NUDX1 gene family. **a** Phylogeny of NUDX1 gene family for sampled *Rosa* genomes. **b** Identified protein domains for NUDX1 gene family by Conserved Domain Database. **c** Gene structure for NUDX1 gene family.

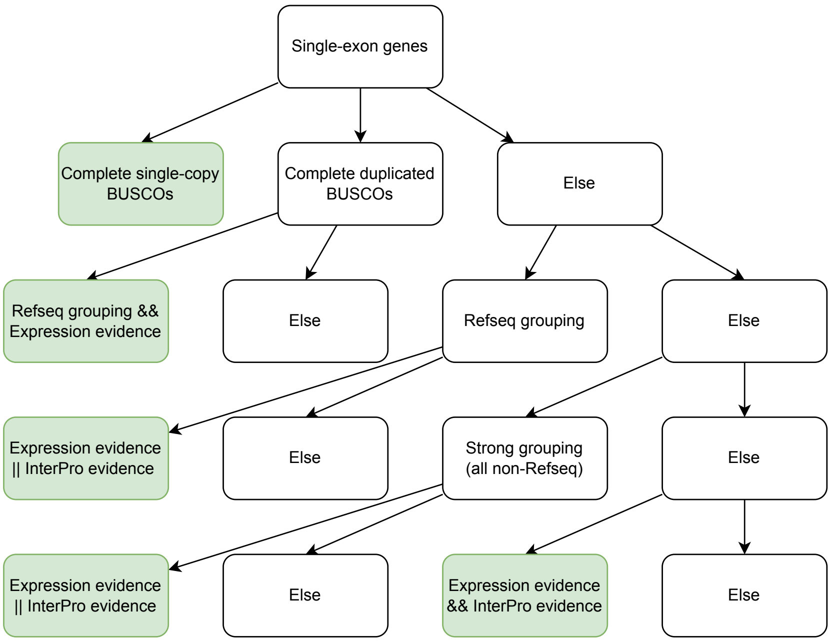


Figure S27: Hierarchical decision tree for single-exon gene filtering. RefSeq grouping: single-exon gene grouped with RefSeq rose annotations; Strong grouping: single-exon gene grouped with all non-RefSeq rose annotations. Single-exon genes fit the criteria in green boxes were kept.
