## Supplemental Materials for "Two high-quality rose genomes underpin a novel *Rosa* pangenome to advance rose genomics, phylogenetics, and breeding"

### Rose pangenome

#### Step 1. Download available rose genomes

##### 1. download Rosa Chinensis (CH) genome

```
wget https://ftp.ncbi.nlm.nih.gov/genomes/all/GCA/041/222/415/GCA_041222415.1_ASM4122241v1/GCA_04122241
wget https://figshare.com/ndownloader/articles/26888665/versions/1
```

##### 2. download Rosa Chinensis (NP) genome

```
wget https://www.rosaceae.org/rosaceae_downloads/Rosa_chinensis/Rchinensis-genome.v1.0/assembly/OBDH_1.
wget https://www.rosaceae.org/rosaceae_downloads/Rosa_chinensis/Rchinensis-genome.v1.0/genes/OBDH_1.0_g
```

##### 3. download Rosa Chinensis (NG) genome

```
wget https://ftp.ncbi.nlm.nih.gov/genomes/all/GCF/002/994/745/GCF_002994745.2_RchiOBHm-V2/GCF_002994745
wget https://ftp.ncbi.nlm.nih.gov/genomes/all/GCF/002/994/745/GCF_002994745.2_RchiOBHm-V2/GCF_002994745
```

##### 4. download Rosa hybrida genome

```
wget https://figshare.com/ndownloader/articles/22774097/versions/1
```

##### 5. download Rosa roxburghii (BMC) genome

```
wget https://download.cncb.ac.cn/gwh/Plants/Rosa_roxburghii_Rr_wild_GWHEROQ00000000/GWHEROQ00000000.gen
wget https://download.cncb.ac.cn/gwh/Plants/Rosa_roxburghii_Rr_wild_GWHEROQ00000000/GWHEROQ00000000.gff
```

##### 6. download Rosa roxburghii (TPJ) genome

```
wget ftp://ftp.cngb.org/pub/CNSA/data4/CNP0004212/CNS0724874/CNA0069570/Rrox.chr.fa.gz
```

##### 7. download Rosa rugosa (HR) genome

```
wget https://www.rosaceae.org/rosaceae_downloads/Rosa_rugosa/Rosa_rugosa_v1.0/assembly/Rosa_rugosa_geno
wget https://www.rosaceae.org/rosaceae_downloads/Rosa_rugosa/Rosa_rugosa_v1.0/genes/Rosa_rugosa_genome.
```

##### 8. download Rosa rugosa (WOR) genome

```
wget https://ftp.ncbi.nlm.nih.gov/genomes/all/GCF/958/449/725/GCF_958449725.1_drRosRugo1.1/GCF_95844972
wget https://ftp.ncbi.nlm.nih.gov/genomes/all/GCF/958/449/725/GCF_958449725.1_drRosRugo1.1/GCF_95844972
```

##### 9. download Rosa rugosa (WUR) genome

```
wget https://www.bioinformatics.nl/pangenomics/data/WUR_RRUG_BELMONTE_hap12.fasta.gz
wget https://www.bioinformatics.nl/pangenomics/data/rrug.hap12.gff3.gz
```

**10. download Rosa sericea (WUR) genome**

```
wget https://www.bioinformatics.nl/pangenomics/data/WUR_RSER_BELMONTE_V1.fasta.gz
wget https://www.bioinformatics.nl/pangenomics/data/rser.final.gff3.gz
```

**11. download Rosa sterilis (TPJ) genome**

```
wget ftp://ftp.cngb.org/pub/CNSA/data4/CNP0004212/CNS0724875/CNA0069571/Rster.newchr.fa.gz
```

The official link to the annotation is currently unavailable. We have created a temporary link.

```
wget https://www.bioinformatics.nl/pangenomics/data/Rster.final.gff3.gz
```

**12. download Rosa wichurana genome**

```
wget https://www.rosaceae.org/rosaceae_downloads/Rosa_wichuraiana/rwichuraiana_BasyesThornless_v1/assem[illegible]
wget https://www.rosaceae.org/rosaceae_downloads/Rosa_wichuraiana/rwichuraiana_BasyesThornless_v1/genes/[illegible]
```

After downloading the genomes, you can use the following command to extract the genome sequences and annotations from the downloaded files:

```
gunzip -c <genome_file.gz>
gunzip -c <annotation_file.gz>
```

#### Step 2. raw data filtering

**1. download pantools-qc-pipeline from PanUtils**

```
git clone https://github.com/PanUtils/pantools-qc-pipeline.git
```

**2. modify the config file `pantools-qc-pipeline/config/config.yaml` to specify the input and output directories and parameters for qc. We applied the following parameters:**

```
min_len: 5000

longest_isoform: True

orf_size: 30
```

**3. run the pipeline**

```
snakemake [rule] --use-conda --cores <threads> [--configfile <config>] [--conda-prefix <prefix>]
```

**4. check the output directory for the filtered data**

#### Step 3. Perform pangenomic analysis using PanTools

Install PanTools according to the instructions (installation through bioconda is recommended). The analysis as described in the manuscript has been performed using PanTools v4.3.1.

**1. build the pangenome using the filtered data**

```
pantools -Xms200g -Xmx400g build_pangenome -t40 <path_to_database> <input_genomes_file>
```

**2. add annotations to the pangenome**

```
pantools -Xms200g -Xmx400g add_annotation -t40 <path_to_database> <input_annotations_file>
```

**3. add phasing information to the pangenome**

```
pantools -Xms200g -Xmx400g add_phasing --assume-unphased <path_to_database> <input_phasing_info_file>
```

**4. perform busco analysis for the pangenome**

```
pantools -Xms200g -Xmx400g busco_protein  <path_to_database> --odb10=eudicots_odb10 -t=40 --phasing
```

**5. perform optimal grouping for the pangenome using busco results**

```
pantools -Xms200g -Xmx400g optimal_grouping <path_to_database> <path_to_database_busco_results> -t=40 -[illegible]
```

**6. choose grouping version based on the optimal grouping**

```
pantools -Xms200g -Xmx400g change_grouping -v=<grouping_version> <path_to_database>
```

**7. perform pangenome classification**

```
pantools -Xms200g -Xmx400g gene_classification \
    --mlsa \
    --phasing \
    --sequence <path_to_database>
```

**8. perform pangenome multiple sequence alignment on subgenome_single_copy_orthologs proteins**

```
pantools -Xms200g -Xmx400g msa \
    -t=40 \
    -H <path_to_database_subgenome_single_copy_orthologs.csv> \
    --align-protein <path_to_database>
```

**9. perform pangenome multiple sequence alignment on subgenome_single_copy_orthologs nucleotides**

```
pantools -Xms200g -Xmx400g msa \
    -t=40 \
    -H <path_to_database_subgenome_single_copy_orthologs.csv> \
    --align-nucleotide \
    --trim-using-proteins <path_to_database>
```

**10. infer a Maximum likelihood phylogeny from SNPs identified from subgenome_single_copy_orthologs**

```
pantools -Xms200g -Xmx400g core_phylogeny \
    -t=40 \
    -H <path_to_database_subgenome_single_copy_orthologs.csv> \
    --align-nucleotide \
    --phasing <path_to_database>
```
